# Engineering *Escherichia coli* Nissle as safe chassis for delivery of therapeutic peptides

**DOI:** 10.1101/2025.08.30.673081

**Authors:** Aaron Pantoja-Angles, Ali Zahir, Sherin Abdelrahman, Cynthia Olivia Baldelamar-Juárez, Shahid Chaudhary, Misjudeen Raji, Luis Fernando Rivera-Serna, Lingyun Zhao, Charlotte A. E. Hauser, Magdy M. Mahfouz

## Abstract

Synthetic biology enables the integration of sophisticated genetic programs into microorganisms, transforming them into potent vehicles for therapeutic applications. Engineering strategies for microorganisms are rapidly evolving, offering promising solutions for cancer therapy, microbiome modulation, digestive health support, and beyond. Developing novel tools to engineer safe, nonpathogenic microbial platforms is essential for advancing clinical therapies. In this work, we present an innovative engineering approach for the probiotic *Escherichia coli* Nissle (EcN), aimed at creating a safe and efficient chassis for the bioproduction of therapeutics. The EcN endogenous pM1 and pM2 plasmids were cured and re-engineered to introduce a CRISPR-Cas12 chromosome shredding device and a therapeutic-producing genetic circuit, thereby generating a nonproliferative therapeutic-delivery system. Next, we build an AI-based bioinformatic pipeline to predict Anticancer-Cell-Penetrating Peptides (ACCPP) candidates. As a proof-of-concept, a selected ACCPP was produced in the engineered EcN chromosome-shredded (CS) chassis. This strategy yields a robust and controllable platform for the safe production and delivery of therapeutics, paving the way for the future development of microbial therapies and their clinical applications.

**GRAPHICAL ABSTRACT:** 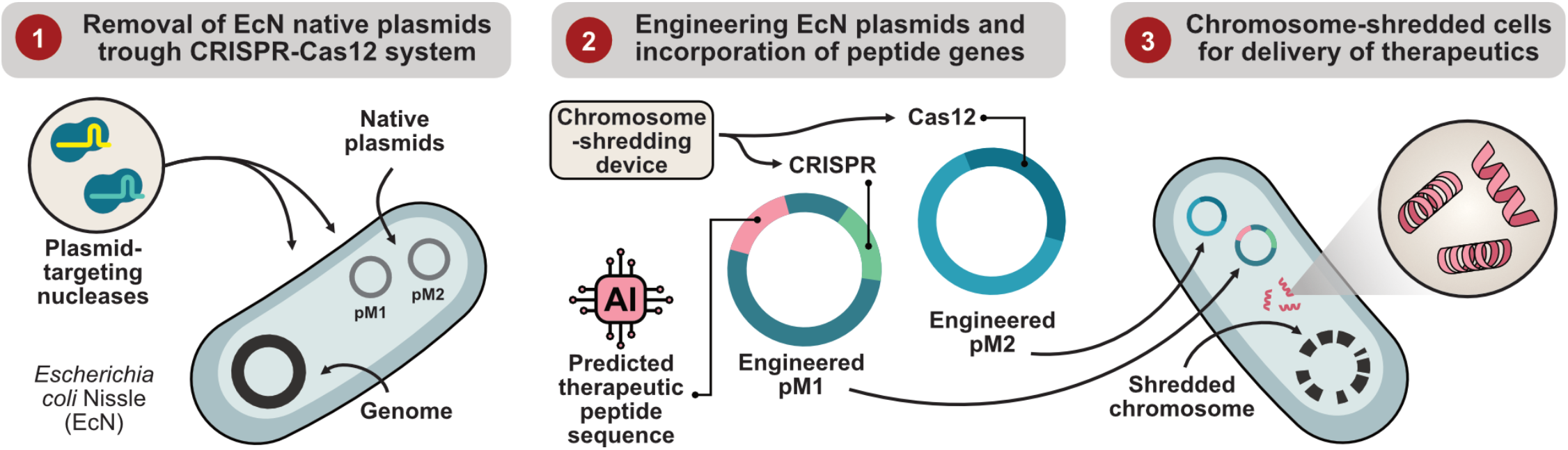

## INTRODUCTION

Advances in synthetic biology are positioning engineered microbes as powerful tools in biotechnology. In recent years, there has been a growing interest in programming bacteria as functional Microbial Therapies (MTs) in the clinical field^1-4^. These living microorganisms offer unique advantages in delivering therapeutic molecules directly to target sites, such as tumors or the gut microbiome ^5, 6^. Moreover, different organisms may selectively target specific tissues, functioning as small biosynthetic factories capable of addressing particular pathologies^7^. Traditionally, MTs have leveraged the inherent properties of microorganisms to combat pathogens and to release anti-inflammatory agents ^8, 9^. Nevertheless, next-generation MTs should be precisely programmed to add extra functionalities^10, 11^, even sense and respond to disease-specific environments^12^, using pathological cues as triggers to activate therapeutic responses^13^.

The development of suitable microbial chassis is a critical step in the creation of effective MTs. An ideal bacterial chassis must withstand the complexities of human microenvironments, avoid triggering harmful immune responses, ensure safety upon administration^14^, and ideally offer proven benefits to the host^15^. One promising candidate for advancing microbial therapies is the *Escherichia coli* Nissle 1917 (EcN) strain^16^. Known for its anti-inflammatory and antimicrobial properties^17-19^, EcN has been particularly valuable in treating gastrointestinal disorders^20^. Additionally, its ability to thrive in hypoxic environments and selectively associate with tumors positions it as a strong candidate for developing novel cancer therapies^21, 22^. Notably, EcN has demonstrated potential for *in situ* production and release of anticancer peptides, making it a promising tool in cancer treatment^8^.

*Escherichia coli* Nissle has further demonstrated its flexibility, as it can adapt its metabolic pathways to utilize unconventional energy sources (such as nitrate^23^, lactate^24-26^, formate^27, 28^, and hydrogen^29^) or to synthesize specific metabolites to contribute to disease mitigation. For example, nitrate is often abundant in hypoxic environments (such as tumors) due to the anaerobic metabolism of cells and tissues^30^. *Escherichia coli*, like other facultative anaerobes, can utilize nitrate as a terminal electron acceptor through anaerobic respiration^31, 32^. Lactate is also commonly elevated in pathological tissues such as tumors^24, 25^ (due to the Warburg effect) or the gut lumen (during inflammation-associated dysbiosis)^28^, which EcN can use as a carbon source^33^. On the other hand, formate can be generated in inflammatory tissues and tumors, and EcN can also use it as an electron donor, making it a useful energy source in these environments. Consequently, EcN is an attractive option for producing more precise disease therapeutic responses.

Generating safe bacterial chassis is essential for potential clinical applications. In addition to selecting nonpathogenic strains, it is important to control their proliferation to ensure that engineered MTs do not outcompete natural microbiota or pose environmental risks after being evacuated from the human body^14^. For instance, safe bacterial platforms are being investigated as bioproduction units with limited proliferative capabilities. Minicells and simcells have been previously described as non-proliferative entities that can express biological products, serving as therapeutics or biosensors for disease detection^34-36^.

We have previously proposed a strategy for generating Chromosome-Shredded cells (CS-cells)^37^, non-proliferative bacterial chassis for safe bioproduction of biologicals. In this work, we further optimized the design and implementation of EcN CS-cells. Our approach begins by using a CRISPR-Cas12-based system to cure the bacterial cells of their endogenous plasmids, pM1 and pM2 (also referred as pMUT1 and pMUT2)^38^. This plasmid-curation step is crucial to eliminate the native plasmids, freeing the cytoplasmic space to introduce engineered versions of pM1 and pM2 that carry the chromosome-shredding device and therapeutic production genetic circuit. pM1 and pM2 are stably maintained across divisions without the stringent requirement for antibiotics, a feature desired in clinical scenarios.

The plasmid-free EcN were transformed with pM1 and pM2, harboring the CRISPR-Cas12 system with different crRNAs targeting the bacterial genome at multiple sites. These crRNAs, in conjunction with the Cas12 nuclease, form ribonucleoprotein complexes that shred the bacterial chromosome, rendering the cells non-proliferative while maintaining functionality for bioproduction capacity, thus providing a versatile and safe framework for microbial therapies. To validate the functionality of EcN CS-cells, we first assessed the hypoxic-mediated expression of GFP, observing that ∼87% of cells actively produced the fluorescent marker, highlighting their sustainability for applications requiring bacterial clearance after therapeutic delivery.

Finally, we envisioned utilizing our developed EcN CS-cells to produce anticancer compounds, focusing specifically on peptides. Peptides, as short molecules, can exhibit diverse properties depending on their amino acid composition^39, 40^, from antimicrobials and antivirals to anticancer. Peptides are highly promising for some applications, while they may have lower affinity, and shorter half-life in comparison to other type of therapeutics, they can be more efficient at self-internalizing into cells and accumulate faster on the targeted tissues^41^. Some efforts have been put into identifying novel anticancer peptides, however several considerations need to be addressed to improve specificity and internalization^42, 43^. To identify novel anticancer peptides with cell-penetrating capabilities, we developed a unique artificial intelligence (AI) pipeline that computationally evolves anticancer peptides and later selects those that may direct their self-internalization into cells. This pipeline enables fast identification of anticancer and cell-penetration peptides (ACCPPs) Using this approach, we selected three peptide candidates that showed efficacy in SW122 colorectal and SHSY5 neuroblastoma cell lines. Finally, we proved as a proof-of-concept the expression of ACCPP2 peptide by our EcN CS-cells, building a robust platform for further characterization as a novel MT.

This study presents an innovative approach for engineering EcN-based CS-cells that combines precision plasmid curing and genome targeting. By leveraging the CRISPR-Cas12 system and the stability of EcN’s endogenous plasmids, we developed a versatile, non-proliferative bacterial chassis capable of carrying previously expressed therapeutics. These advancements lay the groundwork for the future development of microbial chassis, offering promising solutions for *in situ* production and delivery of targeted therapies and overcoming the challenges of safety, specificity, and functionality in clinical settings.

## RESULTS

### A CRISPR-Cas12-based strategy for multiple plasmid curation

To create non-proliferating EcN-MT, we planned to deliver our chromosome-shredding device to EcN using the stably replicating plasmids pM1 and pM2. This can be achieved in two steps. First, the native pM1 and pM2 plasmids must be removed from EcN to allow the transformation of new plasmids with identical origins of replication, preventing plasmid incompatibility. Second, pM1 and pM2 plasmids need to be engineered to carry the chromosome-shredding device (Cas12 and programmed CRISPR array) before being reintroduced into the plasmid-free EcN.

To remove the endogenous plasmids from EcN (a process called plasmid curation), we employed a CRISPR-Cas12-based approach similar to the one used before for chromosome targeting and fragmentation^37^. While previous studies have utilized CRISPR-Cas to remove individual plasmids, our method enables the simultaneous removal of multiple plasmids. (**Fig. 1a**). To achieve this, we introduced two additional plasmids, pCUR1 and pCUR2 (**Fig. 1f**), holding a Cas12 gene and a CRISPR array, both inducible under the presence of arabinose and repressed under glucose. This system produces ribonucleoproteins capable of targeting and clearing all plasmids (pM1, pM2, pCUR1, and pCUR2).

**Fig. 1.**
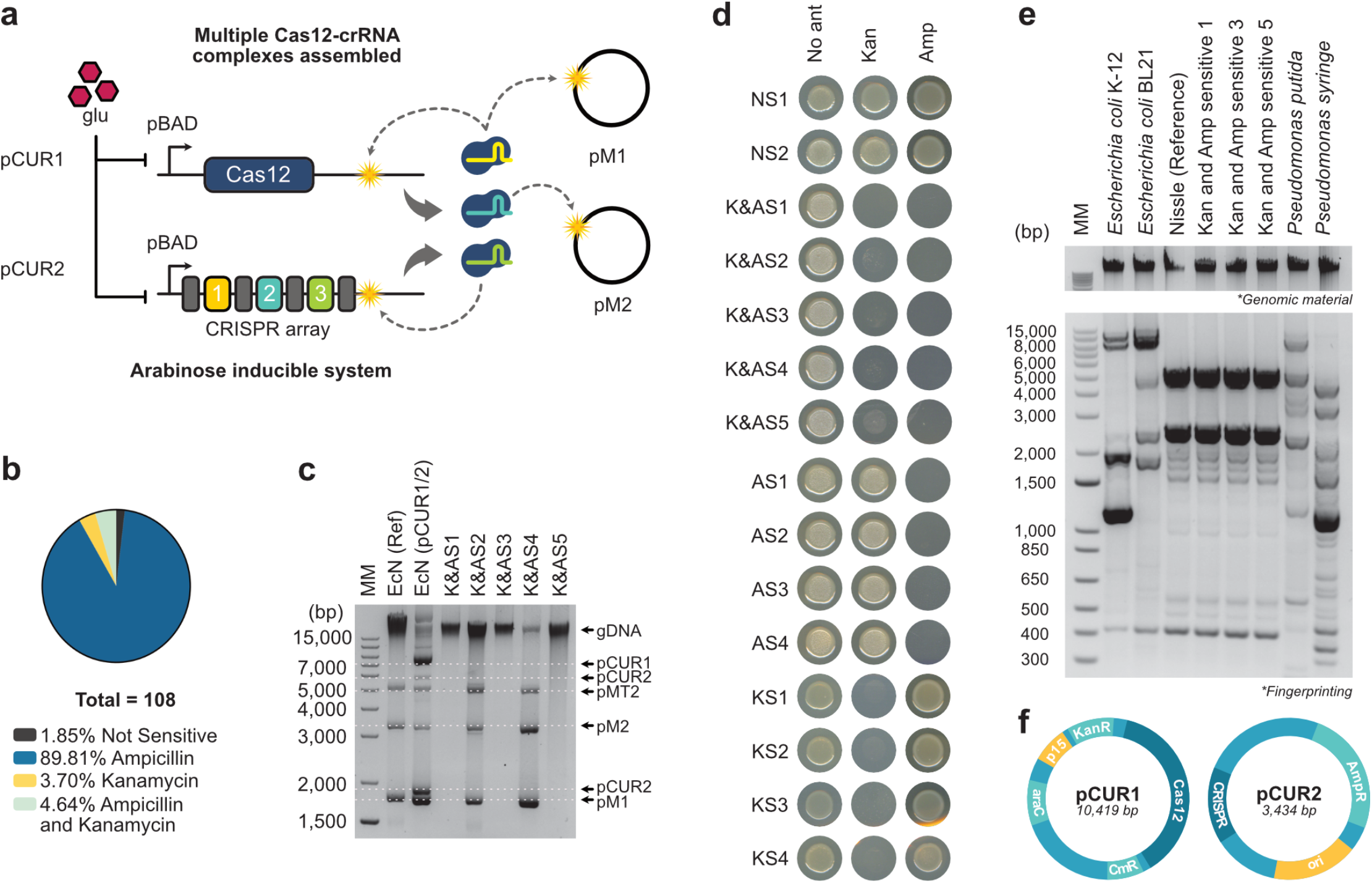
CRISPR-Cas12-based curation of EcN endogenous plasmids. **a.** Schematic of arabinose-induced CRISPR-Cas12 plasmid curation platform. EcN is transformed with pCUR1 (containing Cas12 nuclease) and pCUR2 (containing a CRISPR array with three different spacers). Both Cas12 and CRISPR transcription are repressed under presence of glucose, and activated under arabinose. Upon activation, three different ribonucleoprotein complexes are formed; one leads cleavages on pM1 and pCUR1, another leads the cleavage on pM2 and the third on pCUR2. **b.** Percentage of bacteria showing different sensitivities to antibiotics. Note that 4.64% are sensitive to both ampicillin and kanamycin, suggesting successful elimination of both pCUR1 and pCUR2. **c.** Samples were taken from candidates showing different antibiotic resistance, 2 NS, 5 K&AS, 4 AS and 4 KS. **d.** K&AS candidates were amplified and processed for plasmid extraction together with Reference (Ref) EcN and EcN transformed with both pCUR1 and 2 plasmids (pCUR1/2). Analysis shows successful curation on candidates K&AS 1, 3 and 5. **e.** ERIC1R and ERIC2 primers were used to amplify (ERIC) regions across multiple bacterial species. Top panel is the extracted genomic material used for amplification while lower panel is the fingerprint generated by amplification of ERIC sequences. **f.** Schematics of pCUR1 and pCUR2.

To test the feasibility of this approach, we induced expression of the plasmid-curation system in a triplicate assay (**Supplementary Fig. 1**). From each replicate, colonies (n=36) were recovered and screened for effective plasmid curation. We initially screened for the absence of pCUR1 and pCUR2 plasmids, which carry kanamycin and ampicillin resistance genes. Bacteria sensitive to these antibiotics indicated successful removal of pCUR1, pCUR2, or both pCUR1-pCUR2 plasmids. Samples from each colony were tested for antibiotic sensitivity. Out of all candidates, two colonies showed no antibiotic sensitivity (Not Sensitive, NS), 97 were Ampicillin-only Sensitive (AS), 4 Kanamycin-only Sensitive (KS) and 5 Kanamycin and Ampicillin Sensitive (**Fig. 1b and Supplementary Fig. 2a**). In total, 4.64% of the candidates seemed to have removed pCUR1 and pCUR2, representing an average of 1.6 colonies out of each 36 colonies analyzed (**Supplementary Fig. 2b**).

Next, we selected and extracted the plasmids from all 5 K&AS candidates and from other NS, KS, and AS bacteria to identify their plasmid content (**Fig. 1d**). Thus, we confirmed that the 2 NS bacteria contained all four plasmids as expected (pM1, pM2, pCUR1, and pCUR2), while the KS colonies were missing pCUR1 and the AS colonies were missing pCUR2 (**Supplementary Fig. 3a-c**). Importantly, we show that 3 out of the 5 K&AS colonies contained no plasmid (**Fig. 1c**), demonstrating a curation efficiency of 2.7% for our multiplexed plasmid removal platform. Finally, we performed control experiments to confirm these colonies are indeed EcN and no other microorganism that resulted from the culturing in antibiotic-free conditions. Hence, we compared the genomic fingerprint against the available EcN reference genome. The process relies on the genomic amplification of Enterobacterial Repetitive Intergenic Consensus (ERIC) regions, which have been demonstrated to reveal species and even strain identities^44, 45^. As expected, the curated colonies demonstrated to have a similar pattern as the EcN reference genome but different from other *Escherichia coli* strains or bacterial species (**Fig. 1e**). These results demonstrated the successful application of our developed programmable device for the simultaneous removal of endogenous plasmids from bacterial cells.

### Generation of EcN chromosome-shredded cells

To generate a safe microbial chassis, we integrated an improved version of our formerly described chromosome-shredding device^37^. Leveraging the ability of Cas12 nuclease to self-process their crRNAs, we designed a CRISPR array from which four chromosome-targeting crRNAs can be generated. These crRNAs can increase the number of chromosome-targeted sites as some of them can recognize multiple genomic loci. Furthermore, this principle can be used to design similar chromosome-shredding devices for other organisms (**Supplementary Fig. 4**).

We reasoned that the selection of crRNAs that can target sites on *Escherichia coli* Nissle but not K12 strains could be a proof-of-concept for the specificity of our shredding device. Thus, we created a bioinformatic workflow to eliminate redundant sites present on K12; this strategy can be expanded to eliminate redundant sites present on other species (See **Fig. 2a**); our algorithm returns a list of repetitive sequences (purged dataset) present only in the target genome (EcN) from which a chromosome-shredding device can be generated. From this list, we selected four crRNAs that can target a total of 19 different loci across the EcN chromosome (See **Fig. 2b** and **Extended Supplementary Material 1**).

**Fig. 2.**
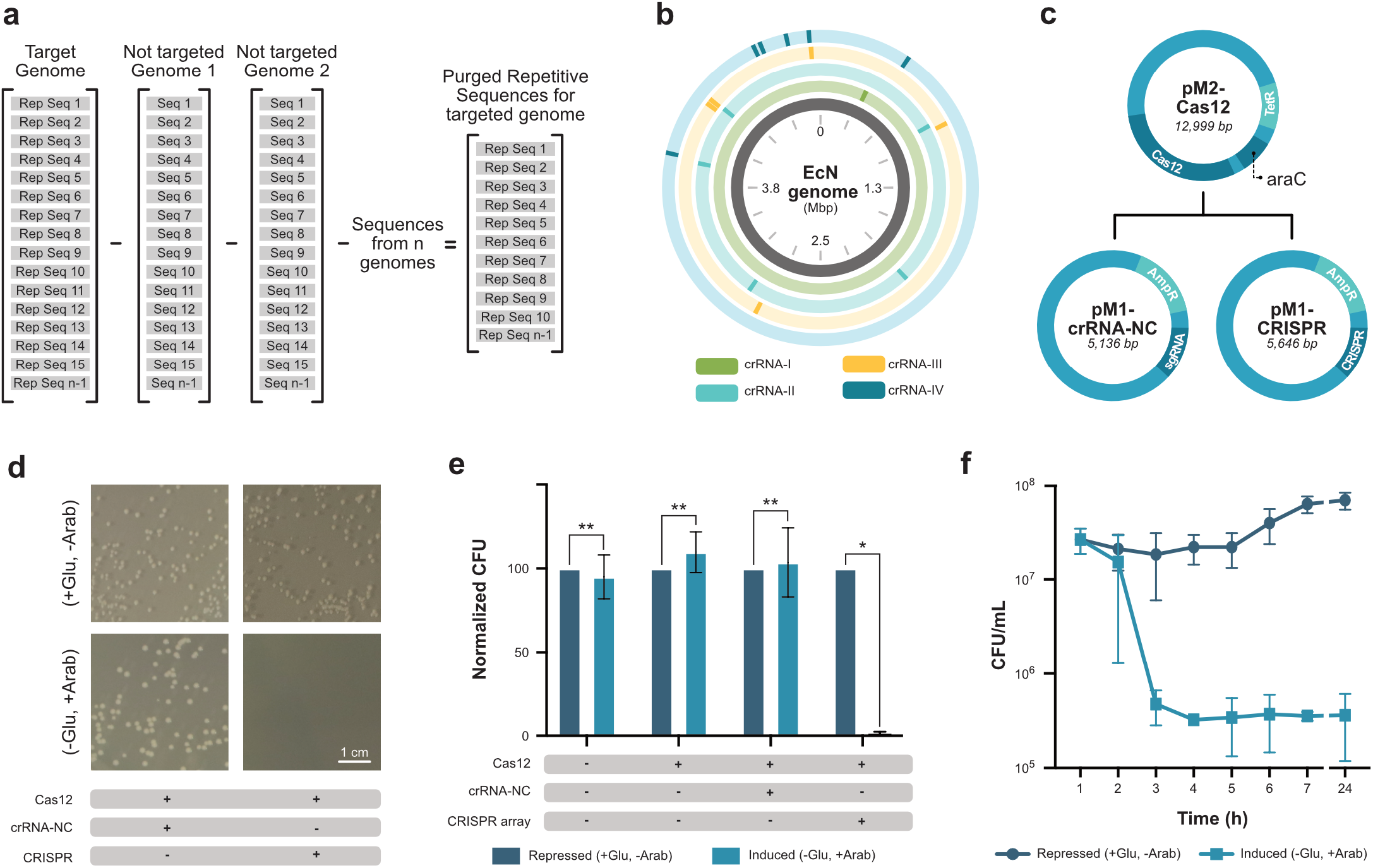
Developing a safe device for generation of EcN CS-cells. **a.** Workflow followed for the selection of EcN targets. After identification of repetitive sequences in EcN genome, identical repetitive sequences found in other genomes are purged. The final list contains repetitive sequences unique to the EcN genome. **b.** EcN genomic representation highlighting the positions at which each crRNA is capable of cleaving. **c.** Schematics of pM1 and pM2 engineered versions containing the chromosome-shredding device **c.** Cas12 and araC genes are contained in pM2. pM1-crRNA-NC version contain a crRNA-NC (not targeting any region of the genome) while pM1-CRISPR harbors spacers giving rise to four chromosome multitargeting crRNAs. **d.** LB agar plates after overnight incubation of EcN variants inducing or repressing the expression of chromosome-non-targeting or chromosome-targeting ribonucleoprotein complexes. **e.** Normalized CFUs of reference EcN or transformed EcN with different plasmid combinations in presence of glucose (glu) or arabinose (ara). Induction of both Cas12 and CRISPR array significantly reduces the proliferation capabilities of EcN. **f.** Plot for numbers of CFU/mL present at each hour after induction of Cas12-crRNA complexes formation. Numbers of CFU/mL at each time is equally calculated for control sample under maintained at a non-permissive state (+glu, - ara). Paired t-test assuming gaussian distribution: * p-value < 0.05 ** p-value > 0.05.

Next, we used engineered pM1 and pM2 versions to deliver our genome shredding device to EcN. These endogenous plasmids have been described as very stable and do not require antibiotic-resistant genes to be propagated. This feature is ideal for their release in humans, as no antibiotics are desired to achieve plasmid stability. As a proof-of-concept, we first utilized pM1 and pM2 versions containing antibiotic-resistant genes to facilitate the cloning process^46^. Hence, we integrated our designed CRISPR array in pM1 (pM1-CRISPR) and the Cas12 and araC genes in pM2 (pM2-Cas12); additionally, we cloned as a negative control a non-targeting crRNA (pM1-crRNA-NC) as shown in **Fig. 2c**. In this sense, pM2-Cas12 plasmid can be combined with pM1-crRNA-NC as a negative control, or pM1-CRISPR for the generation if EcN CS-cells. To ensure tight control over our system, both Cas12 and CRISPR-array are regulated by an arabinose promoter, which ensures a permissive state in the presence of glucose (+glu) and a non-permissive state in the presence of arabinose (+ara).

To characterize the efficiency of our newly designed CRISPR array, we transformed cured-EcN with pM2-Cas12-araC and either pMU1-crRNA-NC or pM1-CRISPR; we hypothesized that activation of our chromosome-shredding platform would result in non-proliferative bacteria. To assess this, transformed EcN cultures were directly plated on either arabinose or glucose agar plates, and colony formation was determined (**Supplementary Fig. 5a, Supplementary Fig. 6, and Fig. 2d**). While single expression of Cas12 or simultaneous Cas12 expression and crRNA-NC transcription did not produce any significant difference in comparison to their repressed reference (+glu, -arab), expression of Cas12 and chromosome-targeting crRNAs produced a significant burden resulting in reduced colony formation. As a result, for every 100 colonies growing in a permissive state, ∼1.6 colonies were proliferating after activating the chromosome-shredding device (**Fig. 2e**). We reasoned that expression and transcription of Cas12 and crRNAs could be performed at lower temperatures at which bacteria do not actively divide, thus avoiding to rapidly dilute the concentration of Cas12-crRNA complexes in each of the daughter cells. Higher concentrations of Cas12-crRNA complexes could result in higher cleavage ratios, hence improving the number of non-proliferative CS-cells. To determine the optimal time of incubation at a lower temperature (16°c), we induced in replicate (n=3) the formation of Cas12-crRNAs complexes and withdrew every hour a small sample that was plated on agar (**Supplementary Fig. 5b**). Next day, colony formation was quantified in every plate to determine the time at which lowest number of escapee bacteria were present. As seen in **Fig. 2f** and **Supplementary Fig. 7**, there is an important inflection point after two hours when a low number of EcN are still capable of proliferating. Particularly after 4 hours, we estimate that ∼1.55 % of the cells are still capable of proliferating, while after 24 hours, the average number of escapees are ∼0.54% (See **Supplementary Table 2**). Therefore, additional methods are necessary to eliminate the escapee cells.

### Purification and test of EcN CS-cell bioproduction capacity

To ensure a pure culture of non-proliferative EcN CS-cells, we treated the cells with ceftriaxone, an antibiotic that inhibits the synthesis of cell wall components. This process selectively targets and lyses proliferating cells, leaving the non-proliferative CS-cells intact. Following ceftriaxone treatment, no proliferative cells were detected in the subsequent culture (**Fig. 3a** and **Supplementary Table 3**). Electron microscopy further confirmed that the morphology of the treated EcN CS-cells remained unaffected (**Fig. 3b**).

**Fig. 3.**
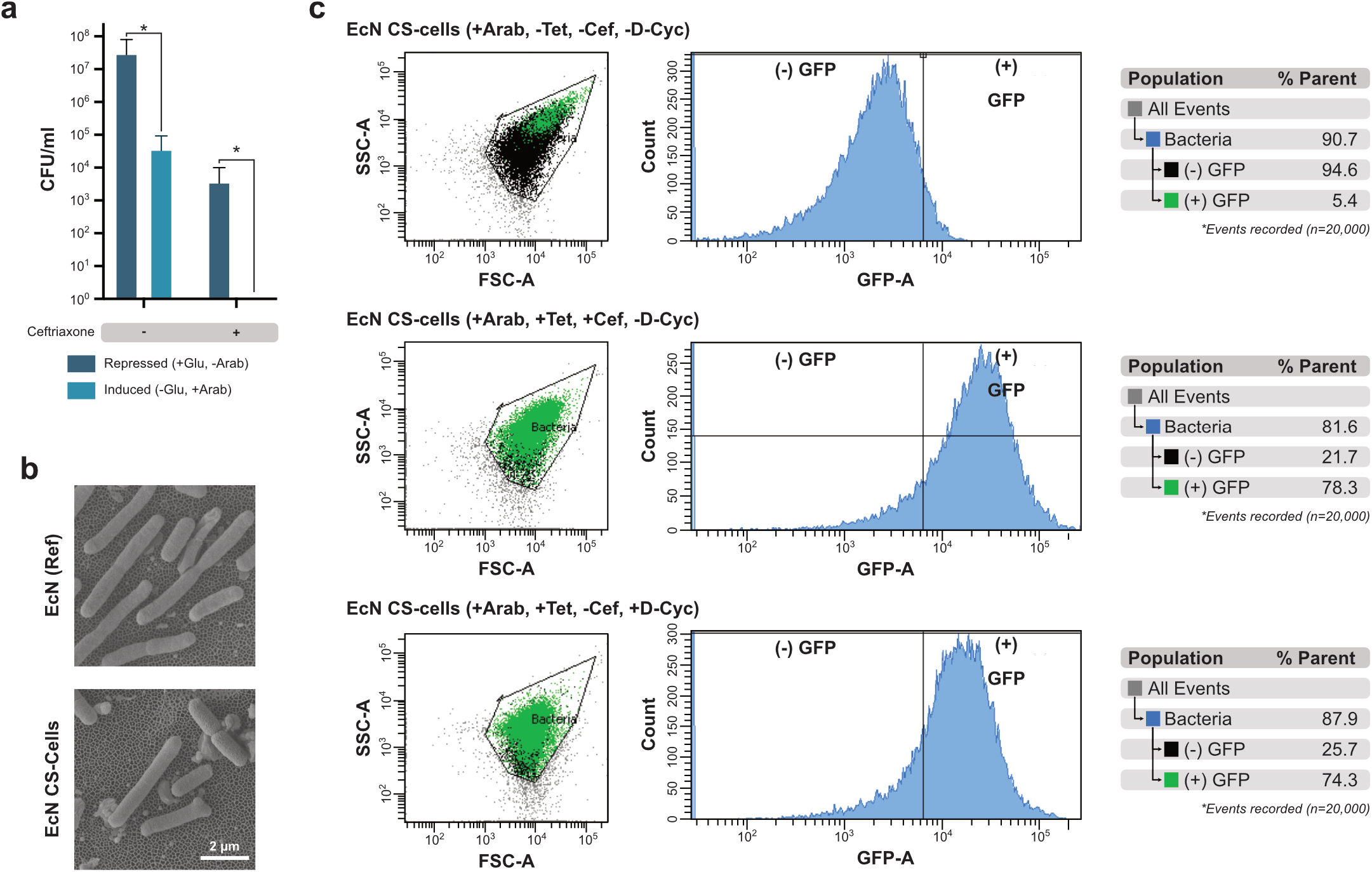
Purification and characterization of EcN CS-cells. **a.** Reference EcN cells and CS-cells are treated with sodium ceftriaxone. Combination of chromosomal-shredding and ceftriaxone leads to undetectable levels of proliferating bacteria. Paired t-test assuming gaussian distribution: * p-value < 0.05. **b.** Electron microscopy on reference EcN and EcN CS-cells. **c.** Flow cytometry plots for Side Scatter (SSC) versus Forward Scater (FSC), histograms on GFP counts and GFP positive/negative quantification. All panels correspond to EcN CS-cells (+Arab; transformed with pM1-CRISPR-Tet-GFP and pM2-C12-araC). Top-panel represent unpurified EcN CS-cells with no GFP induction (-tet). Middle and lower panel are EcN CS-cells with induced GFP expression (+Tet) purified with either ceftriaxone (+Cef) or D-cycloserin (+D-cyc).

We hypothesized that CS-cells could serve as carriers of biological compounds as potential delivery platforms for therapeutic applications. To explore this, we evaluated CS-cells’ ability to express and retain biological cargo after antibiotic-based purification. We constructed pM1-CRISPR-Tet GFP plasmid containing our chromosome-shredding CRISPR sequence and an added tetracycline-inducible GFP reporter gene. This plasmid, in combination with the pM2-Cas12-araC plasmid, was transformed into cured EcN cells. The induction of our chromosome-shredding device leads to the formation of EcN CS-cells that are functional for the expression and loading of GFP.

Subsequent purification of the culture using either ceftriaxone or D-cycloserine ensured that only CS-cells containing the GFP cargo remained. To quantify the efficiency of our process, we employed FACS analysis, which showed that approximately 78.3% of EcN CS-cells displayed GFP fluorescence after ceftriaxone treatment, while 74.3% were fluorescent following D-cycloserine treatment (**Fig. 3c**). This demonstrates that CS-cells can successfully carry and retain biological cargos, highlighting their potential as versatile delivery platforms. Therefore, we reasoned to re-engineer our system to produce novel peptides with the potential for cancer treatment.

### AI-based prediction of new anticancer-cell-penetrating-peptides

We aim to leverage our previously developed bacterial chassis as a delivery vehicle for biologicals, focusing on peptides as therapeutic agents, with a particular interest of those with anticancer properties. Several efforts have been made to identify novel and more efficient anticancer peptides; however, their internalization into cancer cells post-delivery remains challenging. Utilizing Artificial Intelligence (AI) tools has demonstrated an effective tool to predict new therapeutic biologicals with specific properties^47^; we reason to develop a strategy to use these tools in the prediction of peptides with both anticancer and cell-penetrating features.

To predict novel ACCPPs, we relied on previously developed AI modules AntiCP 2.0.^48^ and MLCPP 2.0^49^, which can score input peptide sequences for their potential to be anticancer or cell-penetrating. We first developed a linear bioinformatic pipeline where random peptide sequences of 30 amino acids were analyzed first by the AntiCP 2.0 module, thinking those with scores above 0.9, could be further analyzed by the MLCPP 2.0 module, aiming to identify peptides with high scores on both modules (**Supplementary Fig. 8a**). Nevertheless, after scanning a library of a billion peptides, only 2 of them were given a score higher than 0.9 (**Supplementary Fig. 8b**). This is a bottleneck as we would need to generate thousands of good anticancer candidates that can be used as an input for the cell-penetrating module. To address this issue, we implemented an AI-assisted evolution strategy (**Supplementary Fig. 9**), which, in a matter of minutes can evolve a random peptide into one with high anticancer scores (**Supplementary Fig. 10**).

Using our AI directed-evolution approach, we obtained several candidates and selected the top 10 high scored candidates for both anticancer and cell-penetration (**Fig. 4a** and **Supplementary Table 4**); similar approach was used to devolve the peptides and obtain non ACCPPs sequences (**Supplementary Table 5**). Interestingly, the amino acid composition of our predicted ACCPPs corresponds to those of already existing anticancer and cell penetrating peptides. The high composition of positive amino acid is a feature for directing the peptides to cancer cells, which usually contain a membrane with a negative membrane (**Fig. 4b**). Moreover, presence of non-polar amino acids can also help to mediate the internalization of these peptides bypassing the hydrophobic nature of the cellular membrane. Interestingly, as predicted by AlphaFold, most candidates seem to display alpha-helix as their main secondary structure, a convenient feature commonly found in cell-penetrating and even anticancer peptides. This is because alpha-helices are a structural resource that can be used to form higher homomeric complexes. Indeed, several pore structures composed of alpha-helix peptides have been shown to commonly interfere with or disrupt the cell membrane interface. Additionally, most of the top candidates are predicted to display a good stability on mammalian models, which is partially determined by the predisposition of their amino acid arrangement to be recognized by the proteasome (**Supplementary Fig. 11**).

**Fig. 4.**
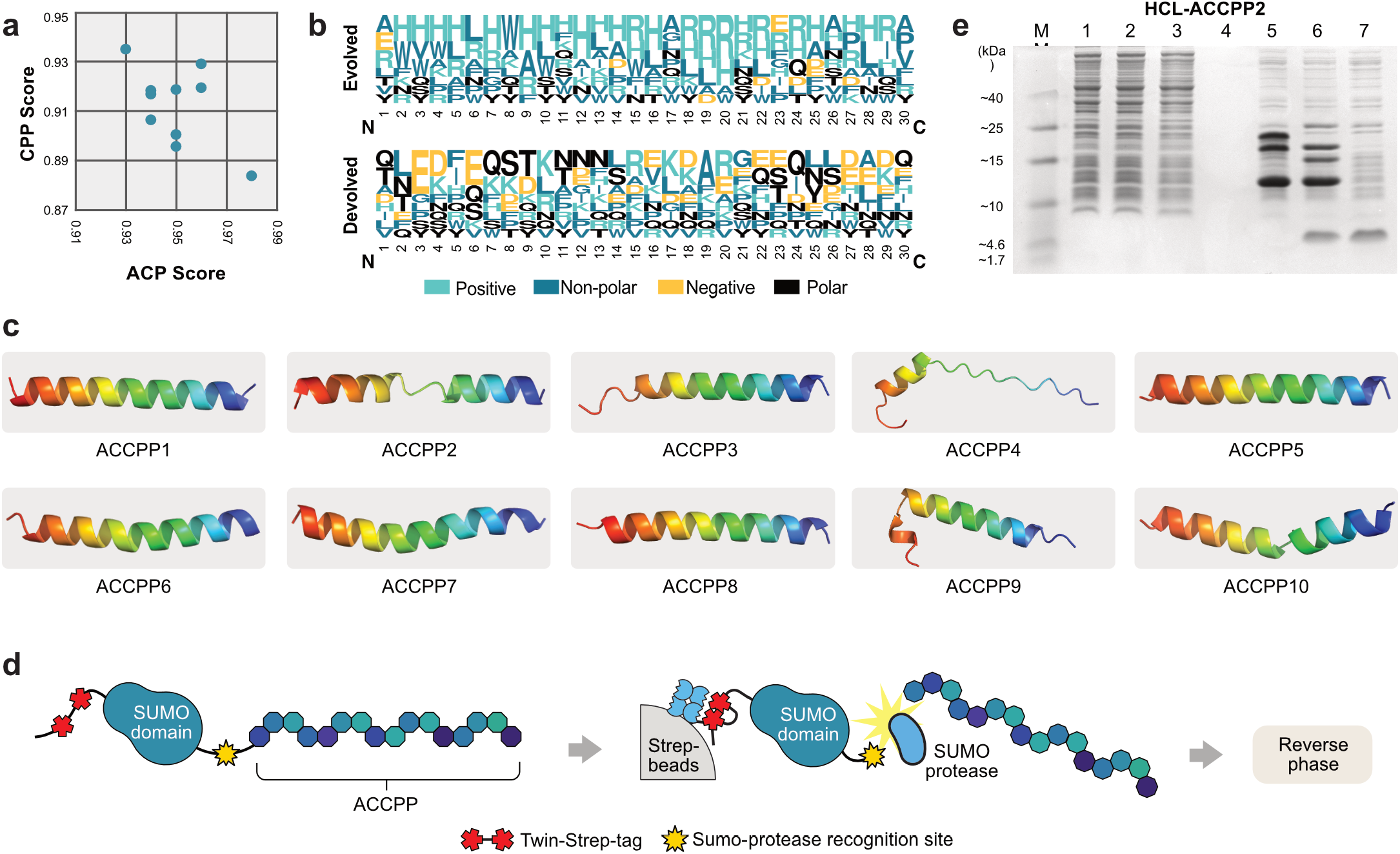
Identification and bacterial expression strategy of ACCPPs. **a.** Scatterplot of final candidate scores (ACP versus CPP), **b.** Amino acid composition of evolved and devolved ACCPPs. Note the composition of hydrophobic, hydrophilic, acidic and base amino acids are enriched differentially. **c.** Alphafold prediction of ACCPP1 candidate. Look Supplementary material for the structural prediction of other peptides. **d.** ACCPP purification strategy based on beads digestion by SUMO protease. SUMO-ACCPP fusion proteins are expressed by bacteria, purified by affinity chromatography, and the peptide cleaved on beads by SUMO protease (Ulp1). Peptide is further purified by reverse phase to remove excess of salts. **e.** Tricine-polyacrylamide gel to run samples from purification process, final recovered protein is shown in lane 7.

We first expressed and purified our peptides from bacteria to test their potential on cancer cells. To do this, we expressed peptides fused to the SUMO domain to support its solubilization (**Fig. 4d**); SUMO-ACCPPs were further purified by affinity chromatography, and peptides were cleaved from SUMO domain on beads by Ulp1 protease (**Fig. 4e**). Peptides were ultimately purified from salt excess by reverse-phase, and purified from endotoxins. While ACCPP1 and ACCPP10 were not successfully cleaved by Ulp1 protease (presumably due to amino acid incompatibilities) the rest of the peptides were purified and assayed. Particularly, ACCPP4 demonstrated to be ∼13 fold stronger on SW122 colorectal cancer cell lines in comparison to Human Dermal Fibroblast (HDF) cells (as non-cancer cells) when treated at 3 µM concentration (**Supplementary Fig. 12a**). ACCPP9 showed around 1.4 fold higher effect on SW122 cells in comparison to HDF (**Supplementary Fig. 12b**). Other peptides did not show a significant higher effect on cancer cells, or showed a lesser effect on cancer cells.

To further indagate the effect of ACCPP4, we tested its effect at the morphological level on different cell lines. While ACCPP4 seem not to have any apparent effect on HDF cells or K562 (lymphoblast), it does affect the morphology of SW122 (colorectal cancer cells) and SH-SY5 (neuroblastoma). Colony formation on SW122 and SH-SY5 is indicative of healthy cell growth, while disaggregating cells from the colony (grape-like structures) are indicative of stress (**Supplementary Fig. 12c**). While further experiments are required, we believe that particularly ACCPP4 could potentially be used in combination with other peptides to boost the effect on cancer cells.

### Generation of CS-cells expressing hypoxic inducible proteins

To produce EcN CS-cells capable of expressing GFP, we incorporated in pM1-CRISPR plasmid a hypoxic-inducible cassette for the expression of either GFP or ACCPP2, as shown in **Fig. 5a**. We selected the pfnrS promoter as it has been shown to be effective for inducing the expression of proteins in hypoxic conditions (like those found in cancer environments). For that, we performed EcN-CS-cell tests on thioglycolate broth, which helped us reduce the oxygen and induce an anaerobic respiration in bacteria. Consequently, low oxygen conditions activate FNR (fumarate-nitrate reduction) protein, which in turn regulates the expression of proteins under the pfnrS promoter^50^.

**Fig. 5.**
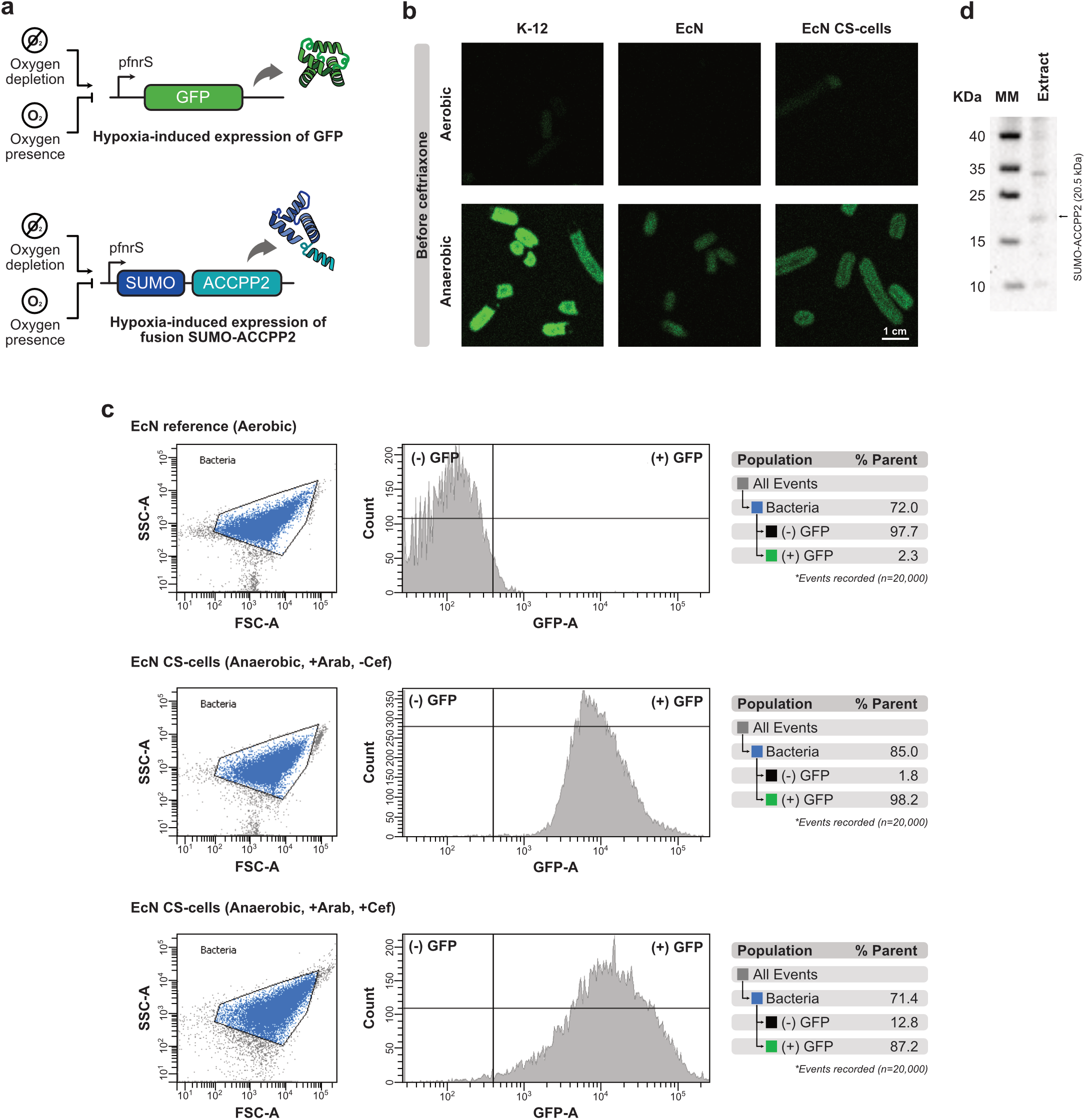
Hypoxic-induced expression of GFP and SUMO-ACCPP2 by EcN CS-cells. **a.** Schematics of genetic circuits for hypoxic induction of GFP or SUMO-ACCPP2 protease. pfnrS promoter is used for the induction of protein expression under anaerobic conditions **b.** K-12 and EcN reference harboring pM1-CRISPR-pfnrS-GFP and pM1-Cas12-araC. EcN is shown before and after conversion into CS-cells. All samples were treated under aerobic and anaerobic conditions to evaluate hypoxic-dependent expression of GFP. **c.** EcN wildtype (grown in aerobic conditions) and EcN CS-cells (grown under anaerobic conditions) before and after ceftriaxone treatments are analyzed trough flow cytometry to determine expression levels of GFP. **d**. Western blot revealing the hypoxic-induced expression of SUMO-ACCPP2 by EcN CS-cells.

To test the effectiveness of this promoter on EcN CS-cells, we first relied on testing the expression of GFP. As a control, K-12 and EcN reference are robustly expressing GFP in the anaerobic conditions, but not in aerobic conditions, as shown by fluorescence microscopy (**Fig. 5b**). Similarly, our EcN CS-cells display robust expression of GFP under anaerobic conditions. Moreover, as shown by flow cytometry, GFP expression is conserved as a cargo in our EcN CS-cells after ceftriaxone purification (**Fig. 5c**).

Finally, to demonstrate that EcN are functional for the expression of our previously predicted ACCPPs, we tested the expression of ACCPP2 under pfnrS promoter. This system is activated in our developed EcN CS-cells under anaerobic conditions. As shown in **Fig. 5d** and **Supplementary Fig. 13**, the western blot reveals the successful expression of SUMO-ACCPP2 by EcN chromosome-shredded cells. This demonstrates the potential of the developed bacterial chassis for holding therapeutic cargoes and for future testing of their functionality as delivery units to targeted organs and tissues.

## DISCUSSION

We have established a non-proliferating bacterial system compatible with therapeutic applications. Our platform demonstrates safety for therapeutic use -due to its non-proliferative nature -and also serves as an attractive option for expressing and storing therapeutic cargoes for potential delivery to the human body. Utilizing CS-cells as cargo carriers would allow the quantification of proteins and peptides stored in CS-cells before administration, potentially leading to more precise doses of delivered therapeutics.

We also considered the advantage of a bacterial system that respond to signals on targeted tissues, therefore, we tested the functionality of a hypoxic-responsive genetic circuit for the expression of biologicals. Successful expression of biologics from hypoxia responsive genetic units demonstrated a sensitive and targeted bacterial chassis for low-oxygen conditions found in tumor microenvironments^51^. Our EcN CS-cells could also be programmed to target other disease cues by designing genetic circuits responsive to cellular states such as oxidative stress^52^, pH changes, or inflammatory signals such as histamines^53^.

Our chromosome-shredding device is highly reprogrammable, making it easy to be employed a variety of non-bacterial microorganisms. This is an advantage in comparison to other platforms where chromosome-degradation is limited by nucleases recognizing specific bacterial genomic sequences^34^. This capability is particularly valuable as non-model microorganisms continue to emerge as promising therapeutic platforms, necessitating novel engineering tools to harness their potential effectively.

Further, in contrast to other CRISPR-Cas plasmid curation strategies, our strategy enables the clearance of several plasmids simultaneously. This could facilitate the engineering of model and non-model bacterial strains that naturally may simultaneously possess a range of plasmids (Ti and pAt found in *Agrobacterium tumefaciens^54^*; p42a, p42, p42c, p42d, p42e and p42f found in *Rhizobium etli^55^*; pWW0, pMG1, pS10, PUM505 found in *Pseudomonas spp^56^*). Our approach demonstrated to remove both pM1 and pM2 in a single step, unlike previous methods^46, 57^.

For practical application of our designed synthetic approach, we next emphasize on the potential use of our AI-assisted peptide selection process, which we envision being used to detect peptides with multiple desirable properties (like stability or selective cytotoxicity). By implementing a similar pipeline, various modules can simultaneously assess whether changes in amino acids improve a peptide’s performance for all desired traits. For the current identified candidate peptides, we believe that combining them all together as a cocktail and optimizing their concentrations could enhance their effectiveness against cancer cells without damaging effects on healthy cells. Even though ACCPP4 demonstrated robust activity on SW122 cancer cell line, further investigation is required to determine an effective dosage of the selected peptides for effective treatment.

Our proof-of-concept-design of the CRIPSR-array successfully led to the generation of EcN CS-cells, however, these crRNAs were also able to target *Escherichia coli* K12 strain when our CRISPR-Cas system was transformed on it. To minimize unintended effects on non-target microbiota, the CRISPR array cold be designed by considering the off-target effects on non-targeted genomes. Leveraging microbial pan-genomes as reference points could help avoid targeting a selected range of microorganisms, particularly if horizontal transfer of the device occurs. Even if such transfer were to remain effective in non-target organisms, the resulting inhibition of proliferation would inherently limit further propagation of the chromosome-shredding cassette.

Our system has demonstrated to be a safe platform for the robust expression of proteins. Compared to other systems such as simcells^34^, most of our CS-cells remain biosynthetically active, supporting the production of biological compounds while minimizing the risk of unintended effects. Our innovative approach of engineering of EcN-based CS-cells that combines precision plasmid curing, genome shredding and tissue micro-environment responsive therapeutic production, enabled successfully a non-proliferative bacterial chassis for practical applications. These advancements lay the groundwork for the future development of microbial chassis, offering promising solutions for *in situ* production and delivery of targeted therapies and overcoming the challenges of safety, specificity, and functionality in clinical settings.

## METHODS

### Plasmid curation

*Escherichia coli* Nissle cells were used for consecutively transformed with pCUR1 and pCUR2 plasmids. EcN containing the pCUR1 plasmid was cultured in presence of 0.5% glucose and 100 µg/mL kanamycin, while EcN containing the pCUR2 plasmid was cultured in the presence of 30 µg/mL chloramphenicol. Single colonies of EcN containing both pCUR1 and pCUR2 were used to inoculate 1 mL of 2xYT media supplemented with 10 mM arabinose and no antibiotics. Cultures were incubated overnight at 16 °C and 180 rpm to induce Cas12 expression and trigger the simultaneous removal of endogenous plasmids pM1, pM2, pCUR1 and pCUR2. After overnight incubation, cultures were serially diluted at a 1:1,000 ratio. From each sample and dilution, 50 µL were plated on LB agar plates without antibiotics. Different dilutions were used to obtain plates from which single colonies could be selected for further screening.

### EcN screening and determination of successful plasmid curation

A total of 108 bacterial colonies (obtained after following the plasmid curation protocol) were picked to inoculate 50 µl of 2xYT medium. Cells were allowed to grow for 4 h at 37°C and 180 rpm. Next, 5 µL from each culture were spotted onto agar plates containing either kanamycin, ampicillin or no antibiotic. Plates were incubated overnight at 37°C. The following day, colonies that grew on plates without antibiotics but not on kanamycin or ampicillin plates were selected to assess the curation of the pM1 and pM2 plasmids. To do this, these colonies were cultured for plasmid extraction using the QIAGEN Maxi-Prep kit. The eluted material was loaded onto gel and quantified by Nanodrop to determine the possible presence of endogenous pM1 and pM2.

### Genomic fingerprinting

Plasmid-cured EcN candidates were analyzed using genomic fingerprinting to confirm their identity. A modified fingerprinting method from James Versalovic et. al. was used. The primers ERIC1R and ERIC2 (5’-CACTTAGGGGTCCTCGAATGTA-3’ and 5’-AAGTAAGTGACTGGGGTGAGCG-3’, respectively) were used to amplify genomic material obtained from *Escherichia coli* bacterial cells. Genomic DNA was extracted from a reference Nissle bacteria, K-12, BL21 and cured cells using the GenElute Bacterial Genomic DNA kit from Sigma Aldrich. Finally, amplification was performed by using NEB Phusion polymerase (98 °C for 30s, 40 cycles of 98 °C for 10s, 60 °C for 1 min and 72°C for 8 min. Final extension at 72 °C for 15 min). Finally, the amplified material was run on agarose gel to determine the amplicon patterns, which are unique to each strain.

### Plate Assay

Cultures were inoculated in triplicate from glycerol stocks (reference EcN strain, EcN with the pM2-Cas12 plasmid, or EcN with pM2-Cas12 plasmid and the pM1 plasmid containing crRNA-A, crRNA-B, crRNA-C or CRISPR array. All samples were supplemented with 0.5% glucose and corresponding antibiotics (no antibiotics for EcN reference). Cultures were incubated at 37 °C and 180 rpm until reaching the exponential phase. Next, glucose was removed from the cultures by centrifugation for 10 min at 4,000 rpm, followed by resuspension of the pellets in fresh 2xYT medium three times. Each culture was then diluted to 1:100, 1:10,000 and 1:1:1,000,000. For each sample dilution, 100 µL were plated on LB agar to activate their specific components (agar containing antibiotics and 10 mM of arabinose), and 100 µL were plated on LB agar in the repressed state to serve as a control (agar containing corresponding antibiotics and 0.5% glucose). Plates were incubated overnight at 37°C or until colonies formed. Plates were photographed and colonies were counted using ImageJ. For each sample, the plate with the dilution factor that allowed the best quantification of colonies was used to calculate the CFUs/mL in the undiluted culture. Finally, counts were normalized to determine the number of colonies formed under induced Cas12/crRNA conditions per 100 colonies in the non-permissive state.

### Liquid Time Assay

Glycerol stocks of transformed EcN (containing pM1-CRISPR and pM2-Cas12) were used to inoculate 10 mL of 2xYT medium in triplicate (supplemented with 0.5% glucose, 100 µg/mL ampicillin and 10 µg/mL tetracycline). Cultures were incubated overnight (∼15 h) at 37 °C and 180 rpm. The next day, 0.5 mL of each culture was taken and used to inoculate 30 mL of fresh 2xYT media (containing 0.5% glucose, 100 µg/mL ampicillin and 10 µg/mL tetracycline). Cultures were incubated at 37 °C and 180 rpm until reaching a OD_600_ value of 0.5-0.6. Cells were washed to remove glucose by centrifugation for 10 min at 4,000 rpm and resuspension of the pellet in 2xYT medium three times (medium containing ampicillin and tetracycline only). Each replicate was then divided into two 10 mL cultures: one was supplemented with 0.5% glucose (to keep CRISPR/Cas repressed) and the other with 10 mM arabinose (to induce CRISPR/Cas activation). Cultures were incubated at 16°C and 180 rpm for 24 hours. During the first 8 hours and after 24 hours, a small sample was taken and diluted at different concentrations (1:1, 1:100 and 1:10,000). From each dilution, 100 µl were plated on LB agar containing only ampicillin and tetracycline. Plates were incubated at 37 °C until colonies were formed. Plates were photographed, and colonies were counted using ImageJ. For each sample, the plate with the dilution factor that allowed the best quantification of colonies was used to calculate CFUs/mL in the undiluted culture.

### Colony count

ImageJ was used to count colonies on plates. First, an image of a plate containing distinguishable colonies was opened. Using the circle tool, the plate area was selected (excluding the edges of the plate). Next, the outside was cleared (Edit tab, “Clear Outside” option), and the image was converted to 16-bit (Image tab → Type → 16-bit). To select the colonies, the image was adjusted (Image tab → Adjust → Threshold). By sliding the bars, the region of the histogram that highlights the colonies was selected, and the threshold was applied. To properly segment any clustered colonies, the image was processed (Process tab → Binary → Watershed). Finally, in the Analyze tab, the number of particles was analyzed. Particle size was set to 20–100, and circularity was set between 0–1.

### Generation of EcN CS-cells and protein expression

EcN bacteria transformed with plasmids carrying the chromosome shredding device were cultured in LB or 2xYT media overnight at 37°C and 180 rpm in the presence of 0.5% glucose and corresponding antibiotics. The next day, 0.5 mL of this culture was used to inoculate 30–50 mL of fresh media also containing 0.5% glucose and antibiotics. Cultures were incubated at 37°C and 180 rpm until reaching an OD_600_ of 0.5-0.6. At this point glucose was removed by washing the bacteria with fresh media; three rounds of centrifugation (5 min 4,000 rpm) and resuspensions were performed. The pellet was resuspended in the same volume as the initial culture. Finally, 10 mM arabinose is added along with the corresponding antibiotics, and cells were incubated at 16 °C and 140 rpm overnight. For GFP expression, tetracycline was added at a concentration of 100 ng/µL; for SUMO-peptide expression, EcN cells was washed twice with thioglycolate broth and cultured in hypoxic conditions.

### EcN purification assay

To remove proliferating escapee cells, EcN CS-cells were treated with 100 µg/mL of sodium ceftriaxone. To test the efficiency, biological replicates were incubated in the presence and absence of sodium ceftriaxone at 16°C and 180 rpm. Next, each sample was diluted to different concentrations (1:1, 1:100, and 1:10,000). Then, 100 µL of each dilution was plated on LB agar containing ampicillin and tetracycline. Plates were incubated overnight at 37 °C and 180 rpm. The following day, plates were photographed, and colonies were quantified using ImageJ from the plates with the optimal dilution factor that allowed single colonies to be visualized and counted. The number of CFUs/mL in the undiluted samples was calculated.

### Hypoxia culture of bacteria

Bacterial cultures were grown in thioglycolate broth containing resazurin (0.5% w/v yeast extract, 1.5% w/v tryptone. 0.55% w/v glucose, 0.25% w/v sodium chloride, 0.05% w/v L-cystine, 0.05% w/v sodium thioglycolate, 0.0001% w/v resazurin). The thioglycolate broth was autoclave-sterilized and sealed immediately avoiding agitation of the medium. Before use, it was place in water bath at 100 °C, boiled for 10 min, and cooled to room temperature. To maintain hypoxic conditions, 15 mL tubes were filled completely with media, closed, and sealed with paraffin. Alternatively, inoculation and cultures were performed in a hypoxic chamber. Importantly, sodium ceftriaxone was added approximately 24 hours after EcN bacteria was cultured in the presence of arabinose. Upon arabinose treatment, EcN CS-cells require time to reach a state in which they stop elongating, this delay is necessary to ensure ceftriaxone does not affect them prematurely,

### Scanning Electron Microscopy

The sample was fixed in 2.5% glutaraldehyde in 0.1 M sodium cacolidate buffer, pH=7.4, at 4°C overnight. Then, osmium tetroxide was used to stain the sample. Next, the sample was dehydrated in ethanol and acetone. Afterward, the sample was dried using a critical point drier. Finally, the sample was coated with 10 nm of Iridium and imaged using the Zeiss Merlin microscope.

### Expression of ACCPPs

BL21 bacteria were used for the expression of fusion SUMO-ACCPP. 40 mL of BL21 preculture were used to inoculate 2 L of fresh LB medium containing the corresponding antibiotics. The culture was incubated at 37°C at 180 rpm until the OD_600_ reached 0.5-0.6. At this point, the culture was cooled, and expression of SUMO-ACCPP was induced with IPTG at a final concentration of 0.5 mM. Then, the culture was incubated at 18°C for ∼16 h and 160 rpm. Cells were harvested and lysed by sonication. Next, SUMO-ACCPP fusion protein was mixed with 2 mL of Strep-Tactin Superflow Plus beads. After 1h of incubation at 4°C with gentle motion, the beads were washed. SUMO domain was cleaved overnight using the Pierce High-Capacity Endotoxin Removal Resins. The absence of endotoxin was quantified using the Pierce Chromogenic Endotoxin Detection Kit. Next, peptides were concentrated by lyophilization and resuspended in 1 mL of ultrapure endotoxin-free water. Salts were removed by reverse-phase chromatography. Final quantification was performed using the Pierce BCA Protein Assay Kit. Peptides were lyophilized and stored at −80°C.

### Cell Culture and Evaluation of Anticancer Peptides on Cellular Metabolic Activity

The SW1222, SH-SY5Y, and HDF cell lines were maintained in Dulbecco’s Modified Eagle Medium supplemented with 10% fetal bovine serum and 1% penicillin-streptomycin. K562 cells were cultured in RPMI-1640 medium with 10% FBS and 1% penicillin-streptomycin. All cultures were incubated at 37 °C in a 5% CO_2_ atmosphere. For cell viability assays, each cell line was seeded at a density of ∼4000 cells per well in 96-well plates. ACCPP4 peptide was added to each culture at 2.6 µM.

Cellular metabolic activity, as a measure of cell viability, was evaluated using the CellTiter-Glo 3D Cell Viability Assay which measures ATP levels. Equal volumes of CellTiter-Glo reagent and cell culture media were added to each well. Samples were mixed by pipetting 8-10 times, followed by incubation at room temperature for 25 minutes. Luminescence was measured using a BMG Labtech plate reader to determine ATP levels, which reflect cellular metabolic activity. The effect of peptides on the metabolic activity of SW1222 cancer cells and normal cells (HDF) was assessed by measuring ATP release after a 24-hour exposure to the peptides at different concentrations (1 µM, 3 µM and 5 µM). For each cell line, a non-treated control was included, and ATP levels in treated cells were compared to those in the control. Cellular morphology was assessed through bright-field microscopy using a ZEISS microscope (ZEISS, Germany). Bright-field imaging provided an overview of the morphological effects of peptide treatment on both cancerous and non-cancerous cells.

## QUANTIFICATION AND STATISTICAL ANALYSIS

All statistical analyses were performed using GraphPad, with details described on each individual figure legend.

## ADDITIONAL RESOURCES

## Code availability

Scripts used to identify genomic repetitive sequences and prediction of ACCPP candidates are available as repositories at https://github.com/anglespantoja. Look for “Repetitive sequences” and “ACCPPs” repositories.

## AUTHORS CONTRIBUTIONS

A.P.A., A.Z. and M.M. conceived and designed the research. A.P.A., A.Z., S.A., C.O.B.J and L.Z. conducted the research. S.A. and C.O.B.J. conducted the cell-culture assays. L.Z. conducted the electron microscopy experiment. L.F.R.S. assisted and provided computational support. M.R and S.C. supported and assisted on the bacterial purification of peptides. C.H., M.M. and A.Z. supervised the research. A.P.A., A.Z. and M.M. wrote and edited the manuscript with input from all the authors.

