## Supplementary_material for "Engineering *Escherichia coli* Nissle as safe chassis for delivery of therapeutic peptides"

for

### SUPPLEMENTARY FIGURES

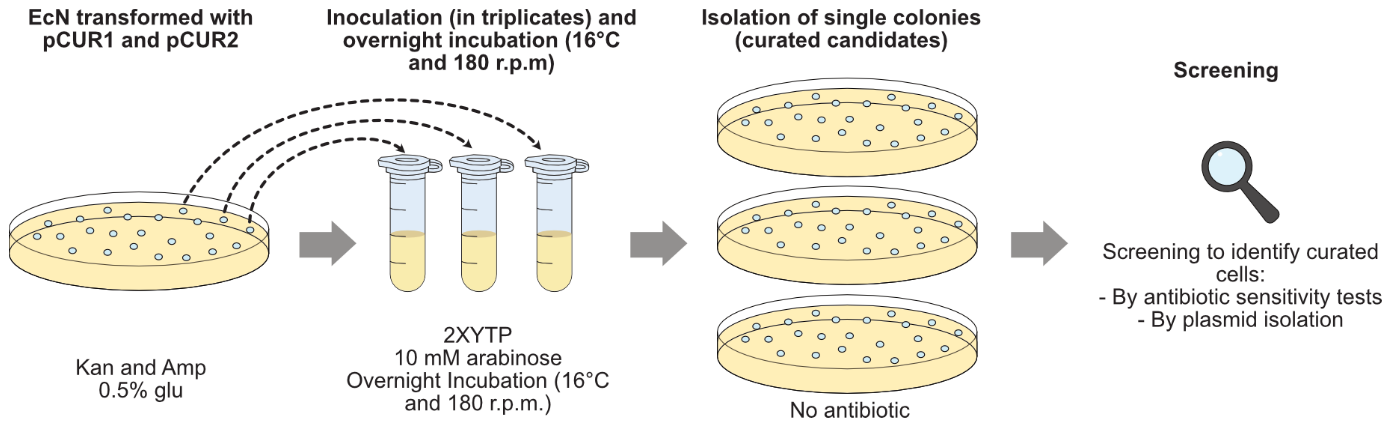

**Supplementary Fig. 1 CRISPR-Cas12-based curation strategy**

EcN cells were transformed with pCUR1 and pCUR2 and maintained under a repressed state (0.5% glucose). Three colonies were picked to perform the curation procedure in triplicates, which consist in incubating the cells overnight at 16°C for expression of the Cas12 and crRNA components (in 10 mM arabinose). Next day, a sample is plated in LB agar plates for colonies to develop. For each replicate, 36 colonies (curated candidates) are taken for further analysis and determination of plasmid curation.

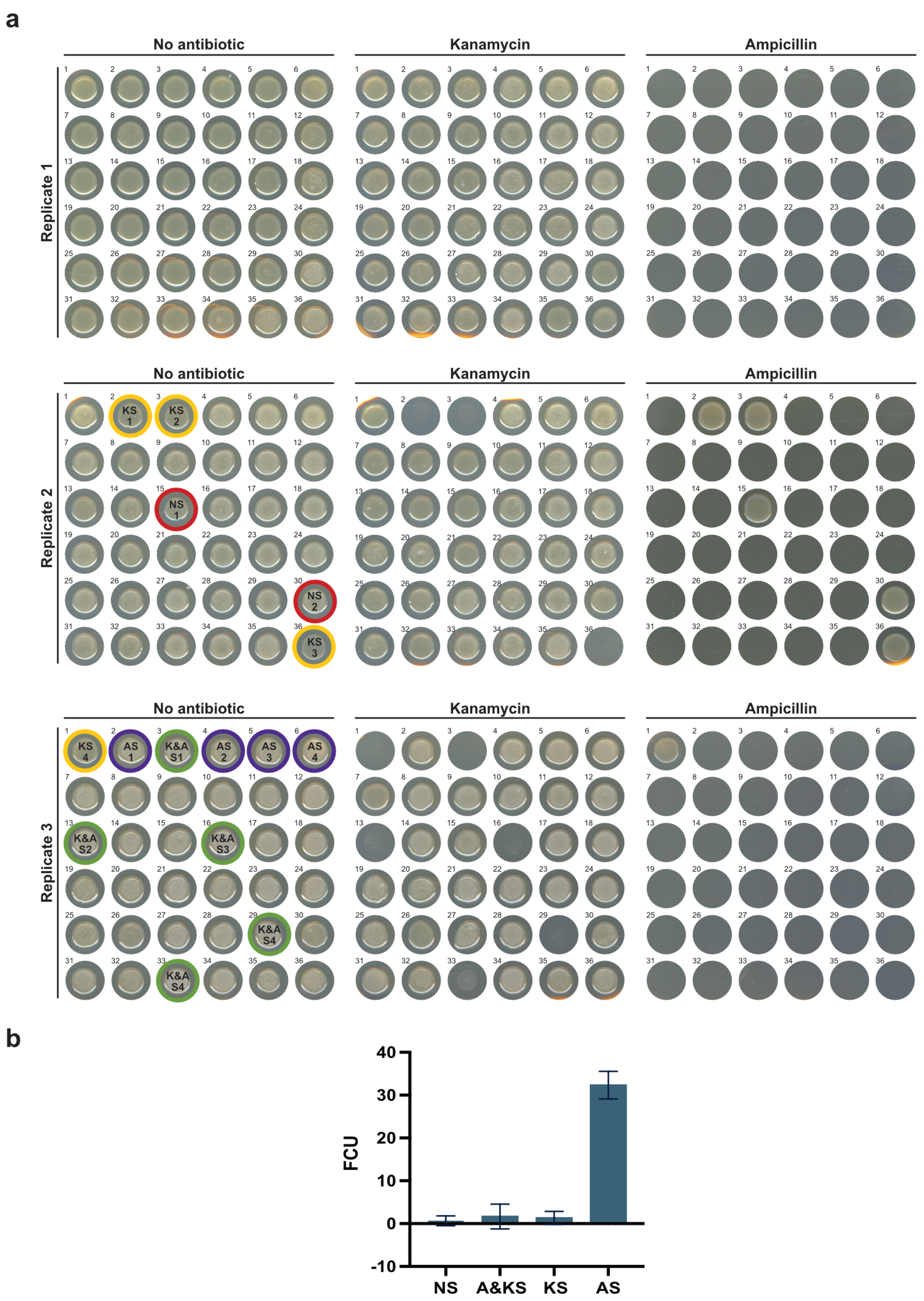

**Supplementary Fig. 2 Screening by antibiotic sensitivity test**

**a.** Antibiotic sensitivity test was used to classify colonies as Non-Sensitive (NS, containing pCUR1 and pCUR2), Ampicillin-only Sensitive (AS, containing only pCUR2), Kanamycin-only Sensitive (KS, containing only pCUR1), or Kanamycin and Ampicillin Sensitive (K&AS, containing no pCUR1 nor pCUR2). Circled colonies were sampled and preserved for further analyses. **b.** Means of colonies obtained across the triplicates displaying different sensitivities to Kanamycin and Ampicillin (NS, AS, KS and K&AS).

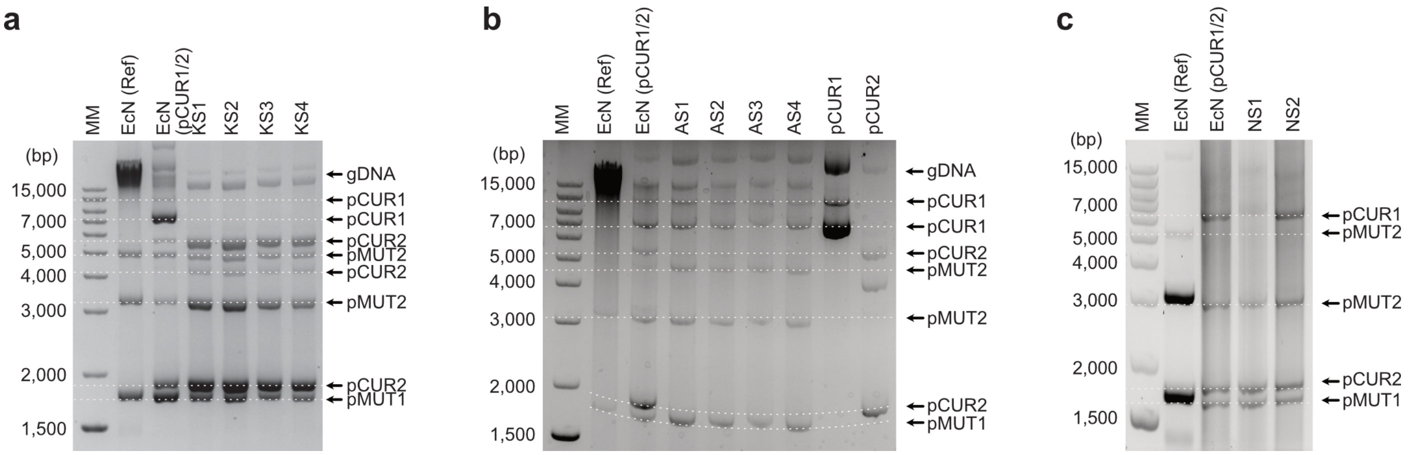

**Supplementary Fig. 3 Extracted plasmids from NS, KS and AS candidates**

**a.** KS colonies lack pCUR1 which contains kanamycin resistant gene, hence its kanamycin sensitivity. **b.** AS candidates lacks pCUR2 which otherwise would confer ampicillin resistance. **c.** NS colonies display all four plasmids (pM1, pM2, pCUR1 and pCUR2). Mind some plasmids are present in two or more conformations.

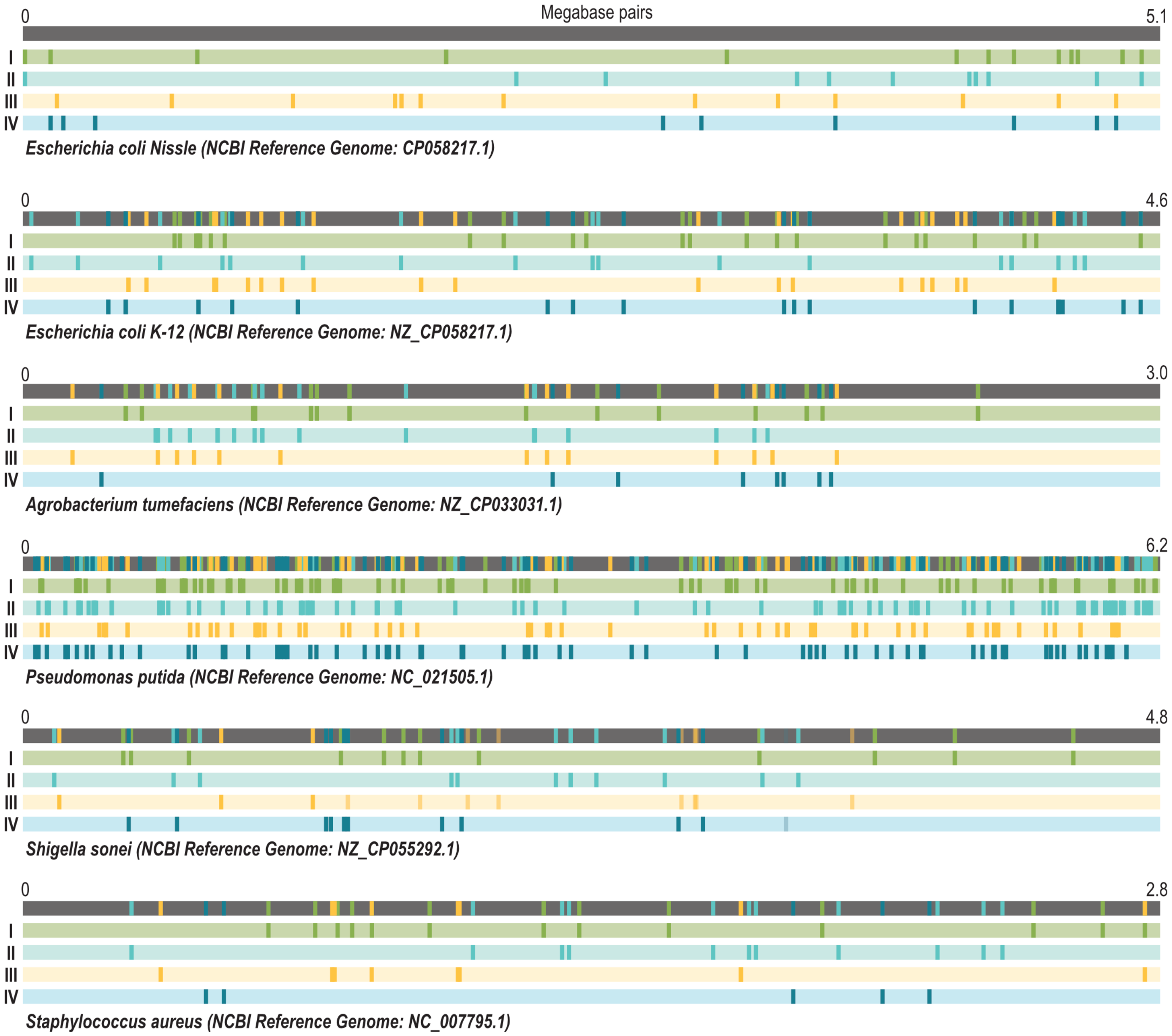

**Supplementary Fig. 4 Schematics of different genomes displaying potential cleavage sites by multitargeting crRNAs**

Four crRNAs are chosen based on their number of potential cleavages and their positions in the genome.

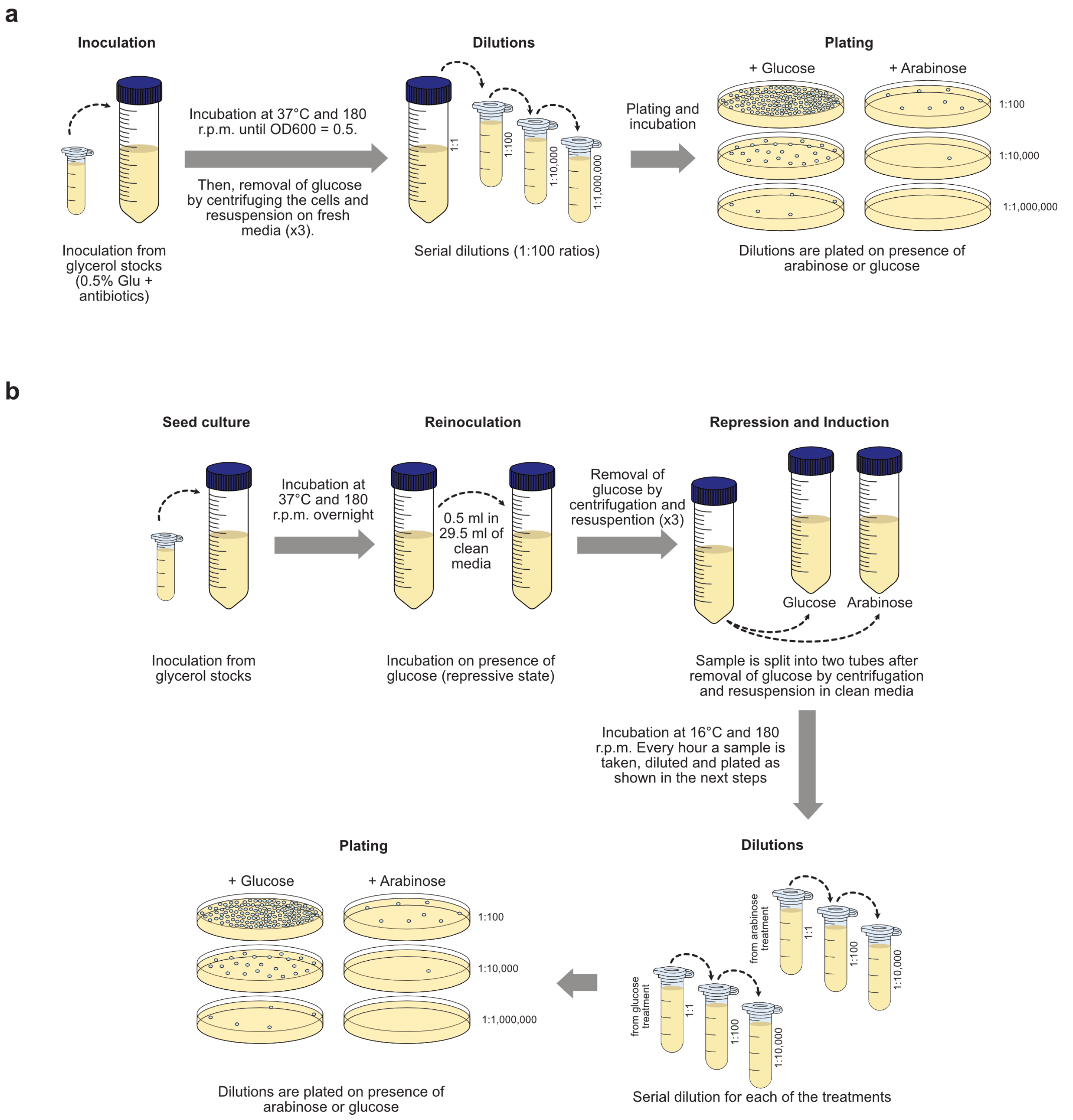

**Supplementary Fig. 5 Experimental procedures for proliferation inhibition assays and optimization of Cas12 expression**
**a.** Experimental procedure to determine the effect of our chromosome-shredding device on EcN proliferation. **b.** Experimental procedure to determine optimal expression and transcription time of Cas12 and crRNAs to achieve highest effect on EcN proliferation.

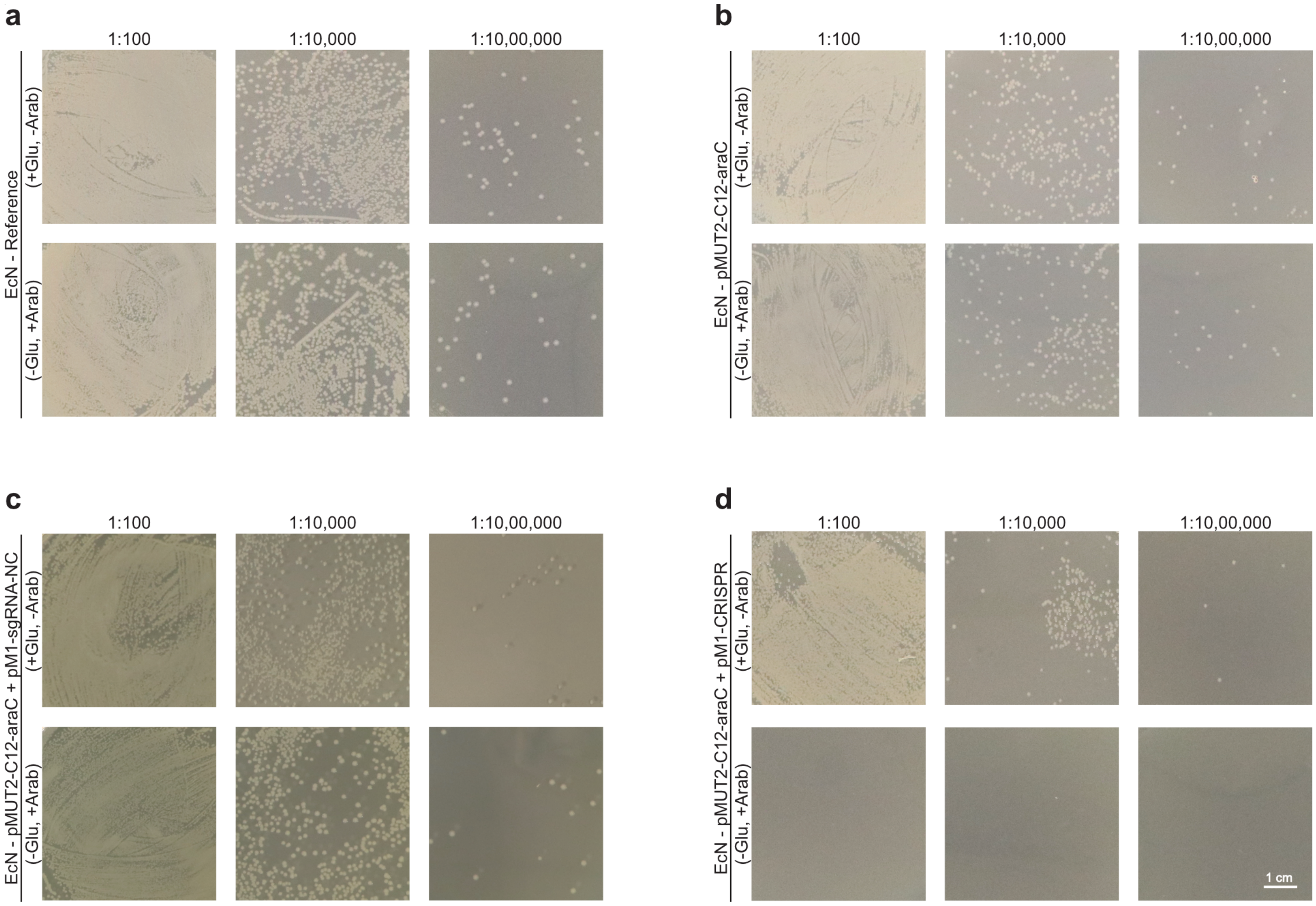

**Supplementary Fig. 6 Determination of bacterial proliferation inhibition**

Different dilutions are plated on agar (1:100, 1:10,000 and 1:10,000,000); dilutions that allow visualization of single colonies are better for determination of proliferation inhibition by comparing chromosome-shredded cells against not chromosome-shredded cells. **a.** EcN cells are used as negative control as they do not contain either Cas12 nor the chromosome-shredding CRISPR array. **b.** EcN cells transformed only with pM-C12-araC is used to evaluate single effect of Cas12, which does not trigger by itself the strong effect on proliferation capabilities. **c.** EcN cells are transformed with both pM2-C12-araC and pM1-sgRNA-NC. Loading of sgRNAs onto Cas nucleases can activate their cleaving effect. However, Cas12 does not produce the expected inhibition in proliferation as the Cas12 nuclease is not being led to their genomic targets. **d.** EcN transformed with pMU12-C12-araC and pM1-CRISPR plasmid can produce a strong inhibition in bacterial proliferation as pM1 plasmid contains the chromosome-shredding cassette. 0.5% glucose represses activation of Cas12 and CRISPR array while 10 mM arabinose induces the expression of this cassette when contained in the cells.

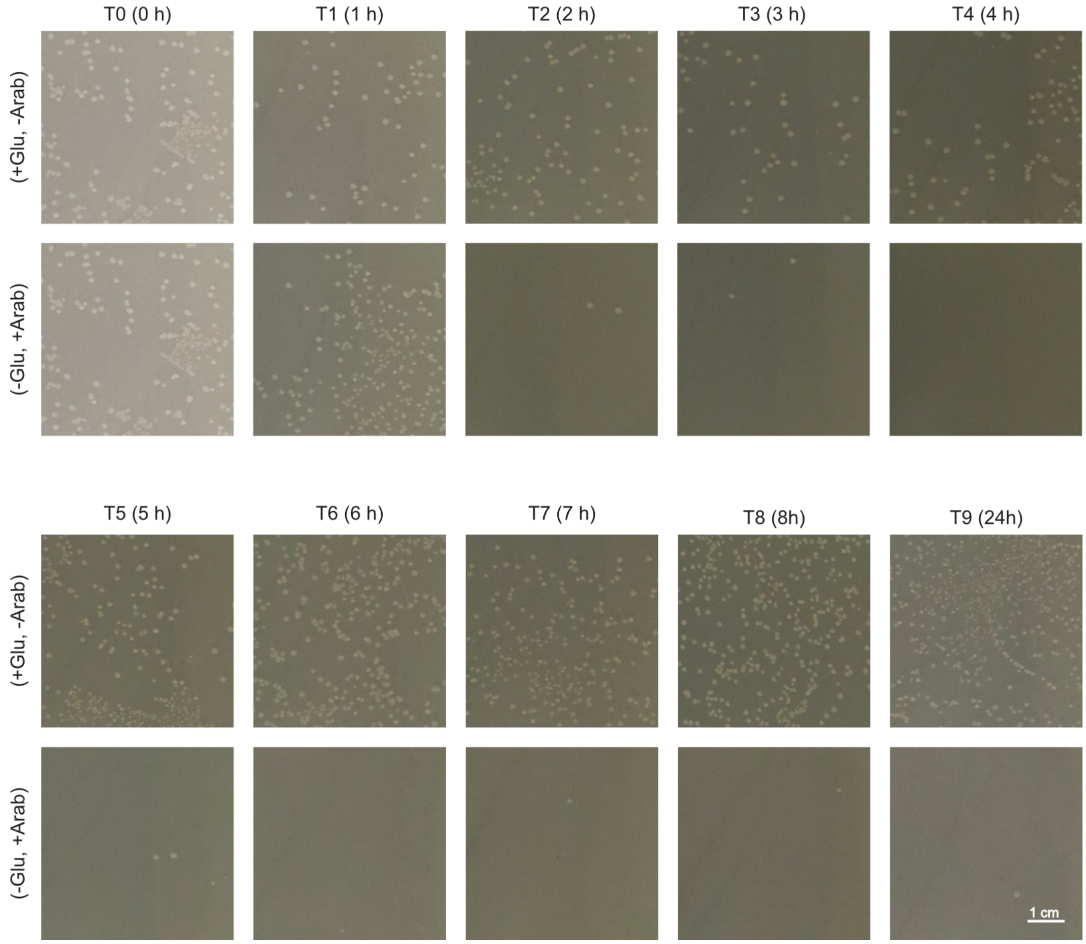

**Supplementary Fig. 7 Determination of effective times for bacterial proliferation inhibition**
The chromosome shredding device is activated upon arabinose induction on EcN cells. Samples after every hour are taken a plated-on LB agar to determine the time with highest effect on bacterial proliferation.

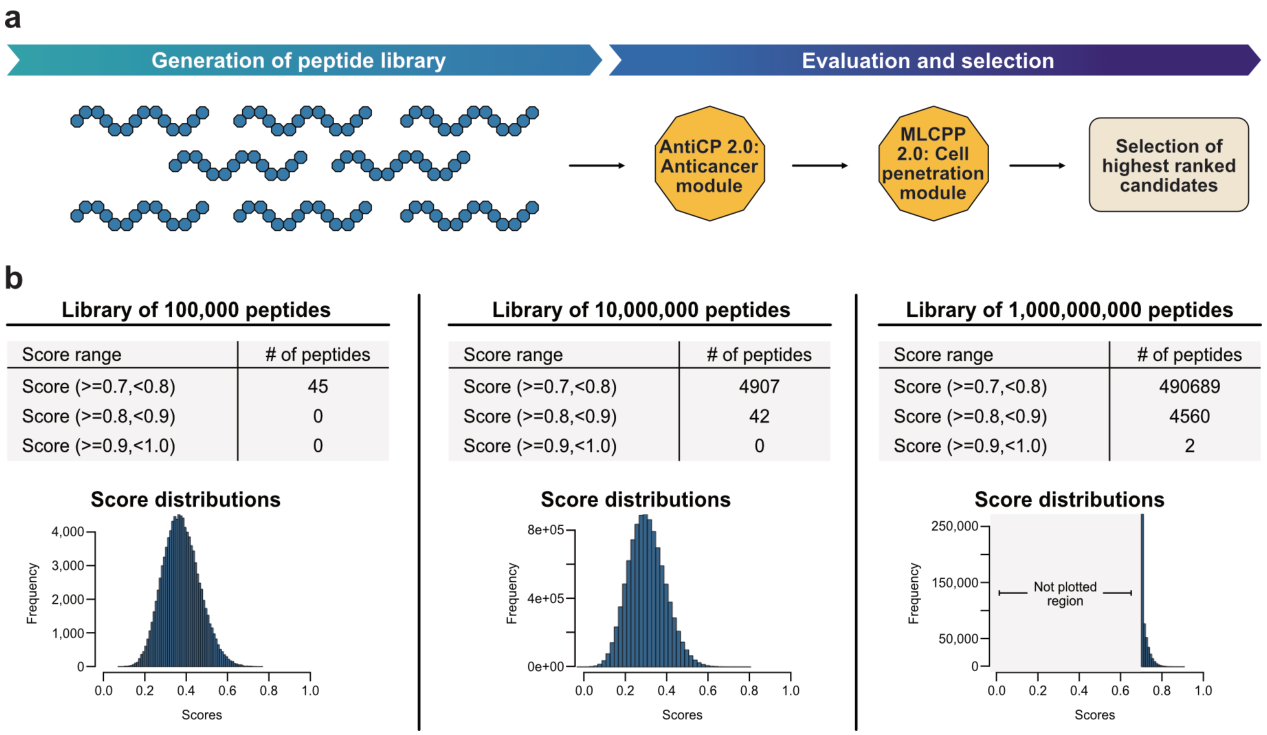

**Supplementary Fig. 8 Bioinformatic pipeline for the identification of ACCPPs and summary of AntiCP 2.0 high-scored peptides**
**a.** A library of random peptides is created, next, they are analyzed by anticancer AI module. Only those with a high anticancer score are consecutively analyzed by the cell penetration module. Finally, highest ranked peptides are selected as potential ACCPPs. **b.** Libraries of randomized peptides of different sizes are run and scored only by AntiCP 2.0 module. Peptides with different scores ranges are quantified and summarized in grey tables. Score distributions are plotted. Only the library of 1,000,000,000 randomized peptide was useful to identify anticancer peptides candidates with scores above 0.9.

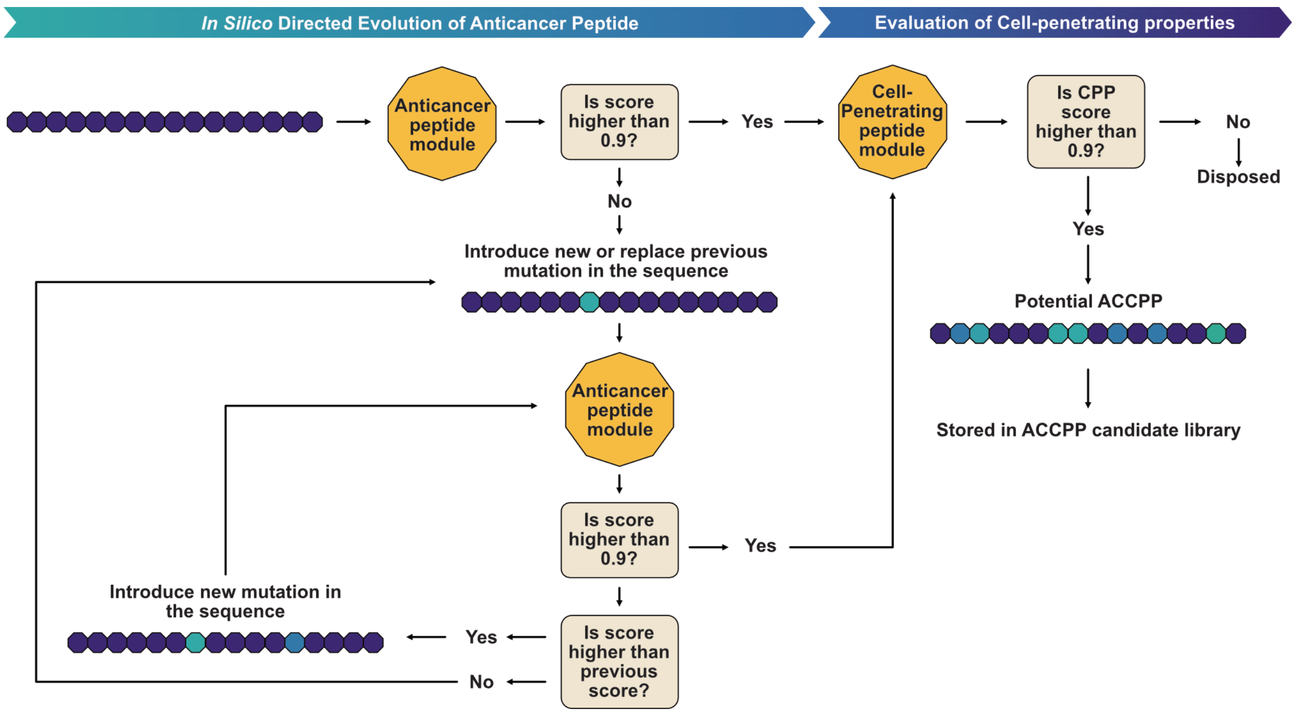

**Supplementary Fig. 9 An AI-assisted approach for in silico evolution of anticancer peptides and identification of ACCPPs**
An artificial intelligence module (AntiCP 2.0) is used for in-silico directed evolution of anticancer peptides. A single random peptide is evaluated trough the module and its anticancer score is calculated, single-amino acids are changed to test its effect on increasing the peptide score. Amino acid substitutions are maintained, removed or added based on its contribution towards increasing the peptide anticancer score. Once the peptide has been evolved, this one is analyzed by the cell penetration module (MLCPP 2.0). If the peptide has a good Cell-Penetrating-Peptide (CPP) score, then it is labelled as an ACCPP candidate, otherwise the sequence is disposed.

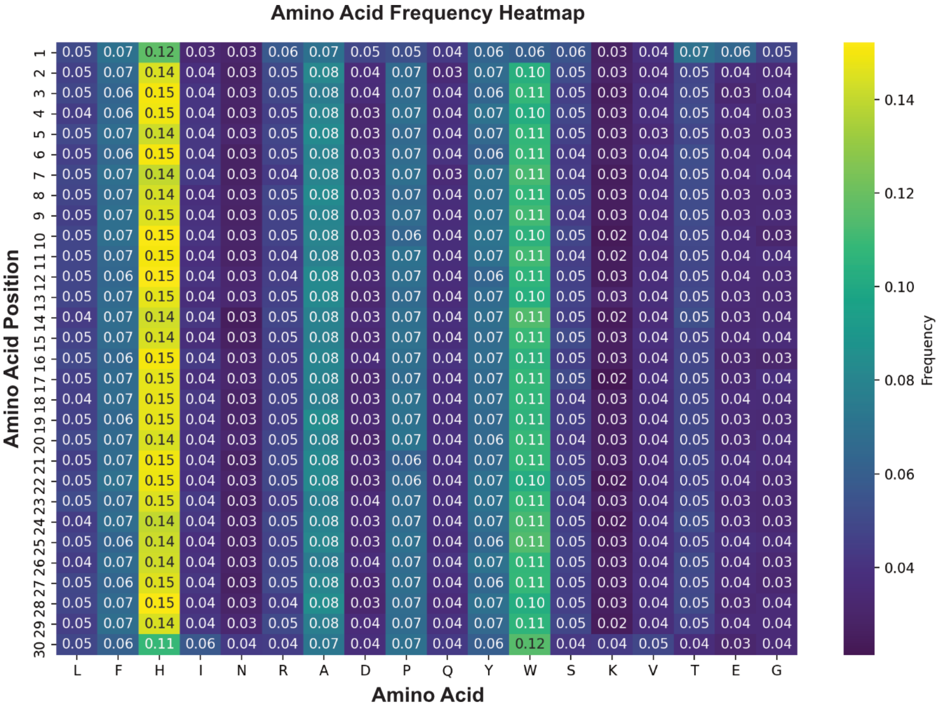

**Supplementary Fig. 10 Amino acid frequency map of evolved anticancer peptides**
Amino acid frequency is calculated for each amino acid position within the evolved peptides (scores above 0.9). Amino acids such as histidine (H) and tryptophan (W) seem to be preferred as part of the composition of the anticancer peptide sequences.

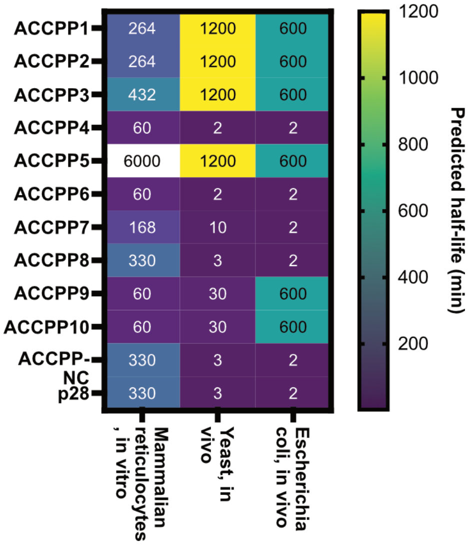

**Supplementary Fig. 11 Half-life estimations for predicted peptides and p28**
Half-life prediction of peptide is calculated using the ProtParam tool. The algorithm analyzes the N-terminal as it is important to determine peptide stability *in vivo*.

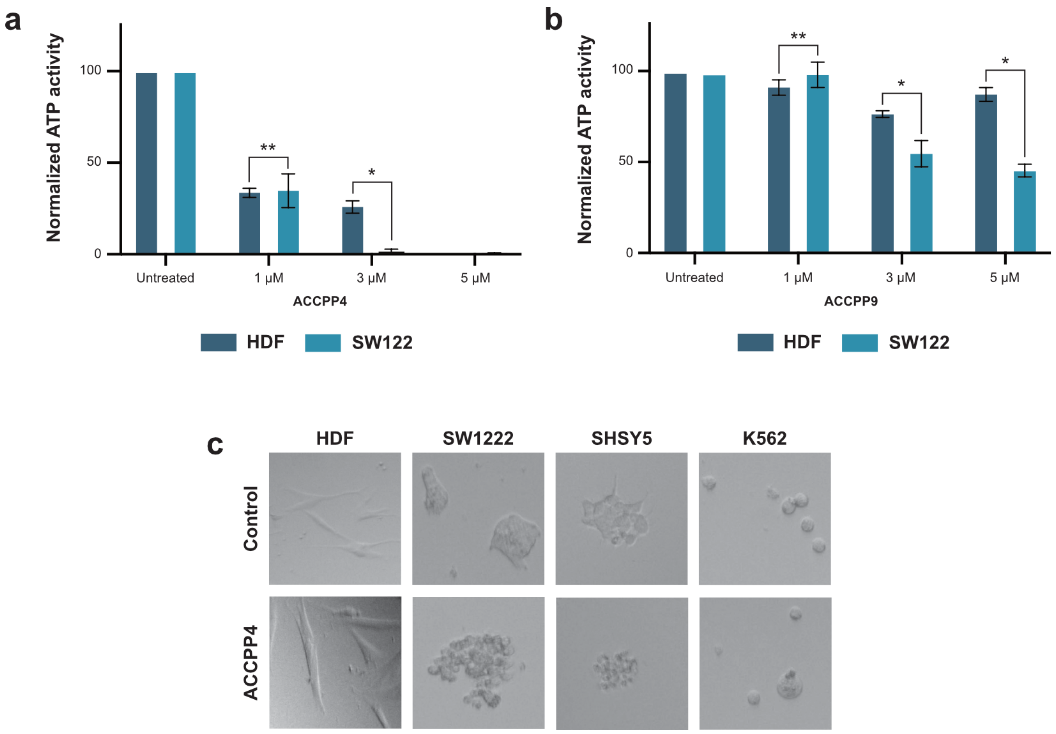

**Supplementary Fig. 12 ATP and cell culture assays to determine the effect of ACCPP candidates**
**a.** ATP-assay is used as an indicative of metabolic activity in HDF and SW122 cells after treatments with ACCPP4 **b.** and ACPP9 **c.** Four different cell lines (HDF, SW122, SHSY5 and K562) untreated and treated with ACCPP4. Unpaired t-test assuming gaussian distribution: * p-value < 0.05, ** p-value > 0.05.

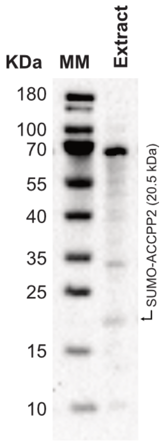

**Supplementary Fig. 13 Immunoblot on EcN CS-cell lysate (extract).**
SUMO-ACCPP2 fusion protein is identified at ~20.5 KDa using an anti-His tag antibody.

### SUPPLEMENTARY TABLES

**Supplementary Table 1 Bacterial colony counts and CFU/mL normalization across samples**

For each bacterial strain, samples under glucose or arabinose were plated at different dilutions. Plates with distinguishable colonies are selected for colony counting. The colony-forming units per milliliter (CFU/mL) were calculated based on the dilution factor of the selected plates. The CFU/mL values for arabinose-treated samples were then normalized against those from their glucose-treated counterparts, serving as the reference condition.

| | | Glucose | | | | Arabinose | | | |
|---|---|---|---|---|---|---|---|---|---|
| | Bacterial strain | Dilution factor | Colonies | CFU/mL | Normalized | Dilution factor | Colonies | CFU/mL | Normalized |
| Replicate 1 | EcN (pM1 + pM2) - Reference strain | 1:1,000,000 | 102 | $1.02x10^9$ | 100 | 1:1,000,000 | 83 | $8.30x10^8$ | 81.37 |
| | EcN (pM2-Cas12-araC) | 1:1,000,000 | 47 | $4.70x10^8$ | 100 | 1:1,000,000 | 45 | $4.50x10^8$ | 95.75 |
| | EcN (pM2-Cas12-araC + pM1-sgRNA-NC) | 1:1,000,000 | 28 | $2.80x10^8$ | 100 | 1:1,000,000 | 26 | $2.60x10^8$ | 92.86 |
| | EcN (pM2-Cas12-araC +pM1-CRISPR) | 1:10,000 | 509 | $5.90x10^7$ | 100 | 1:100 | 10 | $1.00x10^4$ | 1.96 |
| Replicate 2 | EcN (pM1 + pM2) - Reference strain | 1:1,000,000 | 93 | $9.30x10^8$ | 100 | 1:1,000,000 | 100 | $1.00x10^9$ | 107.52 |
| | EcN (pM2-Cas12-araC) | 1:1,000,000 | 19 | $1.90x10^8$ | 100 | 1:1,000,000 | 22 | $2.20x10^8$ | 115.79 |
| | EcN (pM2-Cas12-araC + pM1-sgRNA-NC) | 1:10,000 | 362 | $3.62x10^7$ | 100 | 1:10,000 | 328 | $3.28x10^7$ | 90.61 |
| | EcN (pM2-Cas12-araC +pM1-CRISPR) | 1:10,000 | 419 | $4.19x10^7$ | 100 | 1:100 | 2 | $2.00x10^3$ | 0.48 |
| Replicate 3 | EcN (pM1 + pM2) - Reference strain | 1:1,000,000 | 77 | $7.70x10^8$ | 100 | 1:1,000,000 | 74 | $7.40x10^8$ | 96.10 |
| | EcN (pM2-Cas12-araC) | 1:1,000,000 | 17 | $1.70x10^8$ | 100 | 1:1,000,000 | 20 | $2.10x10^8$ | 117.65 |
| | EcN (pM2-Cas12-araC + pM1-sgRNA-NC) | 1:10,00,000 | 22 | $2.20x10^8$ | 100 | 1:10,00,000 | 28 | $2.80x10^8$ | 127.27 |
| | EcN (pM2-Cas12-araC +pM1-CRISPR) | 1:10,000 | 136 | $1.36x10^7$ | 100 | 1:10,000 | 3 | $3.00x10^3$ | 2.21 |

**Supplementary Table 2 Bacterial colony counts, CFU/mL and calculated percentage of escapees**

Samples were collected every hour from bacterial cultures grown in either glucose or arabinose liquid media. These samples were serially diluted and plated for colony counting. Plates with distinguishable colonies were used to calculate colony-forming units per milliliter (CFU/mL), taking into account the dilution factor. The percentage of escapees was determined by comparing the CFU of cultures grown in arabinose to those grown in glucose. At time zero, CFU counts from glucose and arabinose-treated cells are considered to be the same.

| | | Glucose | | | Arabinose | | | |
|---|---|---|---|---|---|---|---|---|
| | Time (h) | Dilution factor | Colonies | CFU/mL | Dilution factor | Colonies | CFU/mL | % Escapees |
| Replicate 1 | 0 | 1:10,000 | 356 | $3.56x10^7$ | 1:10,000 | 356 | $3.56x10^7$ | 100 |
| | 1 | 1:10,000 | 297 | $2.97x10^7$ | 1:10,000 | 314 | $3.14x10^7$ | 105.72 |
| | 2 | 1:10,000 | 306 | $3.06x10^7$ | 1:100 | 377 | $3.77x10^5$ | 1.23 |
| | 3 | 1:10,000 | 132 | $1.32x10^7$ | 1:100 | 294 | $2.94x10^5$ | 2.23 |
| | 4 | 1:10,000 | 320 | $3.20x10^7$ | 1:100 | 197 | $1.97x10^5$ | 0.62 |
| | 5 | 1:10,000 | 561 | $5.61x10^7$ | 1:100 | 122 | $1.22x10^5$ | 0.22 |
| | 6 | 1:10,000 | 731 | $7.31x10^7$ | 1:100 | 328 | $3.28x10^5$ | 0.45 |
| | 7 | 1:10,000 | 413 | $4.13x10^7$ | 1:100 | 336 | $3.36x10^5$ | 0.81 |
| | 24 | 1:10,000 | 847 | $8.47x10^7$ | 1:100 | 131 | $1.31x10^5$ | 0.15 |
| Replicate 2 | 0 | 1:10,000 | 211 | $2.11x10^7$ | 1:10,000 | 211 | $2.11x10^7$ | 100 |
| | 1 | 1:10,000 | 214 | $2.14x10^7$ | 1:10,000 | 54 | $5.40x10^6$ | 25.23 |
| | 2 | 1:10,000 | 58 | $5.80x10^7$ | 1:100 | 351 | $3.51x10^5$ | 6.05 |
| | 3 | 1:10,000 | 270 | $2.70x10^7$ | 1:100 | 325 | $3.25x10^5$ | 1.20 |
| | 4 | 1:10,000 | 153 | $1.53x10^7$ | 1:100 | 248 | $2.48x10^5$ | 1.62 |
| | 5 | 1:10,000 | 242 | $2.42x10^7$ | 1:100 | 559 | $5.59x10^5$ | 2.31 |
| | 6 | 1:10,000 | 667 | $6.67x10^7$ | 1:100 | 336 | $3.36x10^5$ | 0.50 |
| | 7 | 1:10,000 | 365 | $3.65x10^7$ | 1:100 | 342 | $3.42x10^5$ | 0.94 |
| | 24 | 1:10,000 | 596 | $5.96x10^7$ | 1:100 | 339 | $3.39x10^5$ | 0.57 |
| Replicate 3 | 0 | 1:10,000 | 229 | $2.29x10^7$ | 1:10,000 | 229 | $2.29x10^7$ | 100 |
| | 1 | 1:10,000 | 123 | $2.23x10^7$ | 1:10,000 | 92 | $9.20x10^6$ | 74.80 |
| | 2 | 1:10,000 | 189 | $1.89x10^7$ | 1:100 | 692 | $6.92x10^5$ | 3.66 |
| | 3 | 1:10,000 | 256 | $2.56x10^7$ | 1:100 | 355 | $3.55x10^5$ | 1.39 |
| | 4 | 1:10,000 | 188 | $1.88x10^7$ | 1:100 | 581 | $5.81x10^5$ | 3.09 |
| | 5 | 1:10,000 | 385 | $3.85x10^7$ | 1:100 | 433 | $4.33x10^5$ | 1.12 |
| | 6 | 1:10,000 | 489 | $4.89x10^7$ | 1:100 | 399 | $3.99x10^5$ | 0.82 |
| | 7 | 1:10,000 | 324 | $3.24x10^7$ | 1:100 | 420 | $4.20x10^5$ | 1.30 |
| | 24 | 1:10,000 | 620 | $6.20x10^7$ | 1:100 | 617 | $6.17x10^5$ | 1.00 |

**Supplementary Table 3 Bacterial colony counts before and after treatment with ceftriaxone**

EcN bacterial cells containing pM1-CRISPR and pM2-Cas12-araC were treated both with glucose or arabinose. After repression or induction of the chromosome-shredding device, bacteria were treated with ceftriaxone. Samples were taken before and after ceftriaxone treatments. Samples were serially diluted and plated for colony counting. Plates with distinguishable colonies were used to calculate colony-forming units per milliliter (CFU/mL), taking into account the dilution factor. Colonies were counted and CFU/mL calculated.

| | | (+Glu, -Arab) | | | (-Glu, +Arab) | | |
|---|---|---|---|---|---|---|---|
| | Treatment | Dilution factor | Colonies | CFU/mL | Dilution factor | Colonies | CFU/mL |
| Rep 1 | No ceftriaxone | 1:100 | 75 | $7.50x10^4$ | 1:1 | 227 | $2.27x10^3$ |
| | Ceftriaxone | 1:1 | 0 | 0 | 1:1 | 0 | 0 |
| Rep 2 | No ceftriaxone | 1:10,000 | 32 | $3.20x10^6$ | 1:100 | 6 | $6.00x10^3$ |
| | Ceftriaxone | 1:1 | 0 | 0 | 1:1 | 0 | 0 |
| Rep 3 | No ceftriaxone | 1:10,000 | 872 | $8.72x10^7$ | 1:100 | 101 | $1.01x10^3$ |
| | Ceftriaxone | 1:100 | 11 | $1.10x10^4$ | 1:1 | 0 | 0 |

**Supplementary Table 4 Summary of ACCPP candidates**

Amino acid sequences for the predicted ACCPPs is show. Scores anticancer and cell penetrating scores are shown, as well as the overall ACCPP score.

| Peptide Information | | | ACP - CCP - Scores | | | |
|---|---|---|---|---|---|---|
| ACCPP | Identifier | Amino acid Sequence | ACP Score | CCP Score | CPPE Score | ACCPP Score |
| ACCPP1 | 11767E28I374 | AHWRHLHTASYHWAHLQHRHNRGHETFLHA | 0.96 | 0.9419092 | 0.98633 | 0.944516664 |
| ACCPP2 | 10780E31I271 | AWWARRRQHWKKKAFHPKAWQHRRAVKAWA | 0.96 | 0.9444044 | 0.9736984 | 0.939782525 |
| ACCPP3 | 26E16I269 | TWQHHAHWATHVLHWLRLRRRLPRYHPNIV | 0.95 | 0.9740987 | 0.9431867 | 0.934378456 |
| ACCPP4 | 17253E34I452 | RRHHPPPKAHSHIVHTAYDRRRRQDSPHIP | 0.93 | 0.9627294 | 0.9712696 | 0.932534912 |
| ACCPP5 | 5412E30I588 | VWHWHLHLRYWAADRHLRLRSDERAADIRY | 0.98 | 0.9328975 | 0.9473555 | 0.931892745 |
| ACCPP6 | 3082E22I273 | RKVKFFFWFWHHVHNHGRRHHIFHHWRHLF | 0.94 | 0.9304436 | 0.9868411 | 0.92909997 |
| ACCPP7 | 2404E25I301 | YHSLLLGAHHFHLRRAAHLRKWIQHNHLRR | 0.94 | 0.9390718 | 0.976142 | 0.928333722 |
| ACCPP8 | 7127E12I21204 | LHVHLRHWRLRHHLIRNRHAYHLTFLAHSP | 0.95 | 0.9703866 | 0.9279142 | 0.925217746 |
| ACCPP9 | 15792E24I330 | EFHPAWYWHHVNHHWFWWRRLREDPRRLRL | 0.94 | 0.951501 | 0.9525313 | 0.923167247 |
| ACCPP10 | 16063E29I372 | ENYWRNRYHLQIRPRHVLHHHLHLHAHWQV | 0.95 | 0.9778619 | 0.9160481 | 0.92288428 |
| p28 | Positive Control | LSTAADMQGVVTDGMASGLDKDYLKPDD | 0.26 | 0.9999984 | 0.0015648 | 0.130782375 |

**Supplementary Table 5 Summary of non-ACCPPs (ACCPP-NC)**

Amino acid sequences for the predicted ACCPP-NC is show. Scores anticancer and cell penetrating scores are shown, as well as the overall ACCPP score.

| Peptide Information | | | ACP - CCP - Scores | | | |
|---|---|---|---|---|---|---|
| ACCPP NC | Identifier | Amino acid Sequence | ACP Score | Class | Probability | ACCPP Score |
| ACCPP NC1 | 211E20I164 | LNEYIEKDDLRNNPFEIAAAGQTNYSRELQ | 0.01 | Non-CPP | 0.0330869 | 0.02154345 |
| ACCPP NC2 | 288E14I166 | QQPDWSQPTKTNTSRPGQAREQSQLREAKI | 0.01 | Non-CPP | 0.0338576 | 0.02192878 |
| ACCPP NC3 | 156E15I145 | ITEKRKFEKVYDNQPVLKARWFEQQSHEDY | 0.01 | Non-CPP | 0.0343397 | 0.02216984 |
| ACCPP NC4 | 349E19I200 | DTYDQQSSQNTIIRRRVFPVSYTPLNEQVL | 0.01 | Non-CPP | 0.0380327 | 0.02401634 |
| ACCPP NC5 | 389E17I257 | VLEDFELKYRAKHELVLDAPYEQQNIWTNE | 0.01 | Non-CPP | 0.0405997 | 0.02529986 |
| ACCPP NC6 | 54E21I181 | NLKNEKQPIRQYDANDQYYVSKKVVPLTYE | 0.03 | Non-CPP | 0.0226035 | 0.02630175 |
| ACCPP NC7 | 175E20I213 | DQQQSVPKQKTRVTDNSFIFEARLRHKREA | 0.03 | Non-CPP | 0.02388217 | 0.02694109 |
| ACCPP NC8 | 5E30I177 | QLSSYSQFTKNPFLKLDVRHKNSVFEDAKF | 0.04 | Non-CPP | 0.0147859 | 0.02739295 |
| ACCPP NC9 | 469E21I267 | NELFYSLDLQPGQPLDFITNPFYTNQQFQY | 0.02 | Non-CPP | 0.03557544 | 0.02778772 |
| ACCPP NC10 | 264E21I238 | TFEKHVQSTKNVKSDQKDKKGLRTEGYIFR | 0.01 | Non-CPP | 0.0456937 | 0.02784686 |

### EXTENDED SUPPLEMENTARY MATERIAL 1

Genomic regions are displayed surrounding the cleavage sites targeted by each designed crRNA.

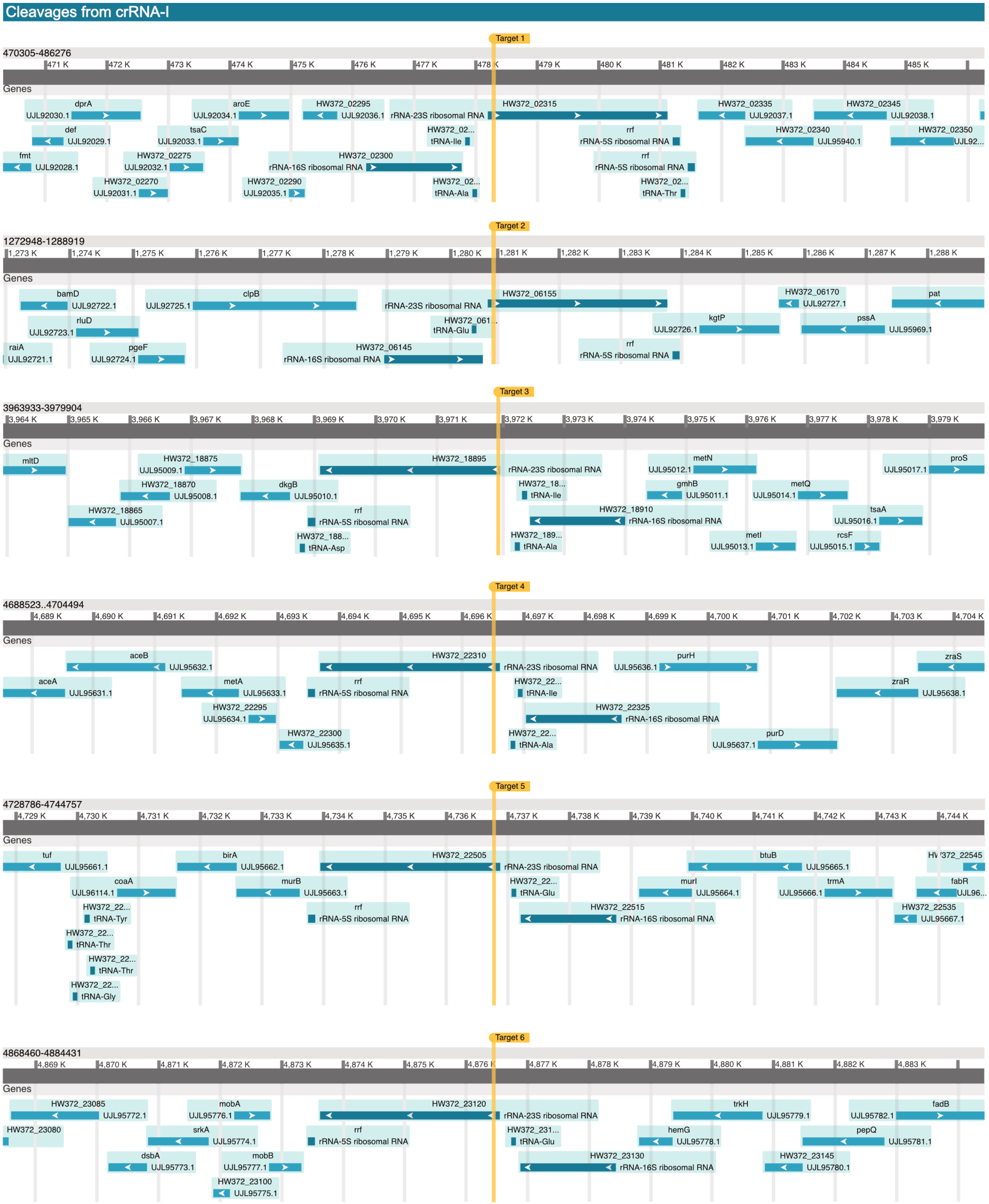

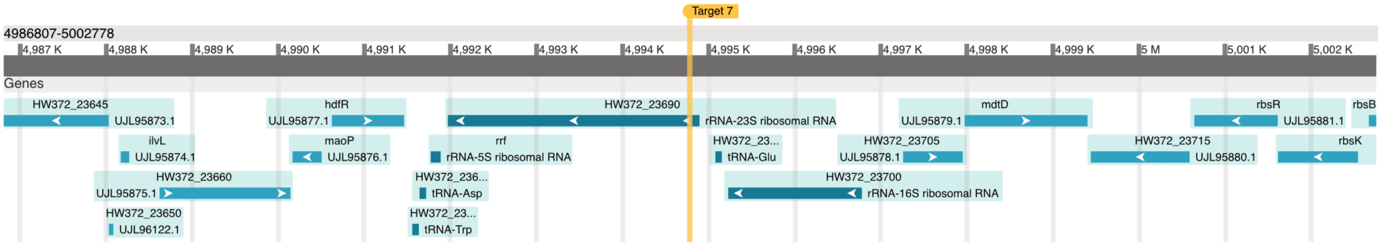

### Cleavages from crRNA-II

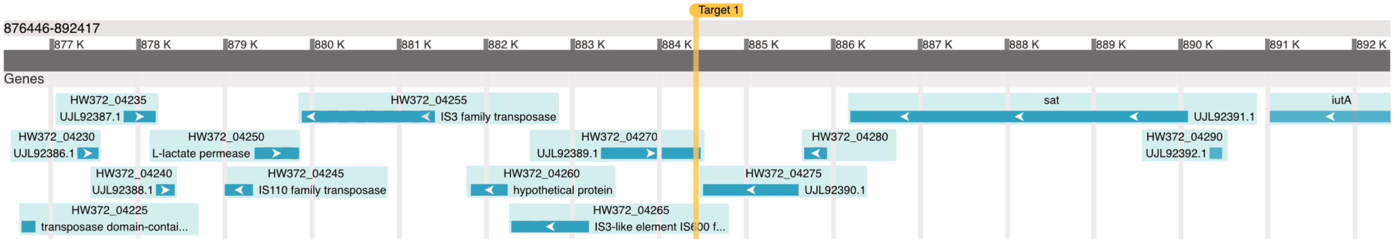

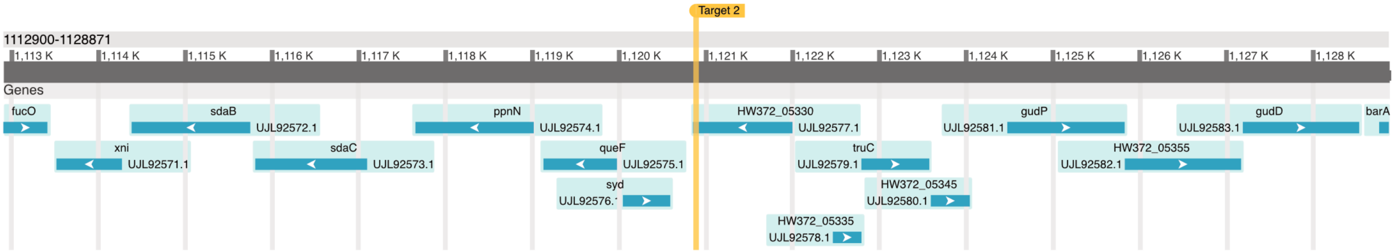

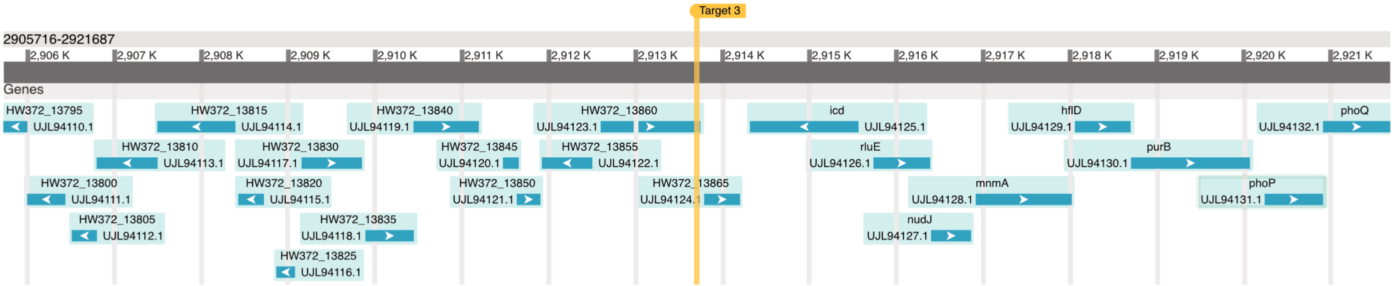

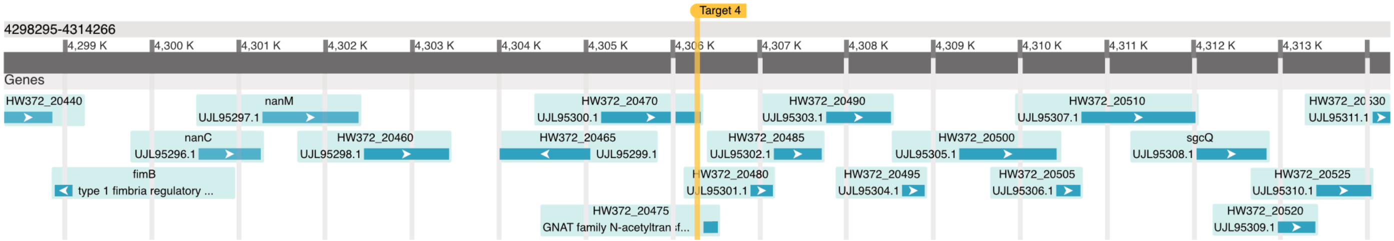

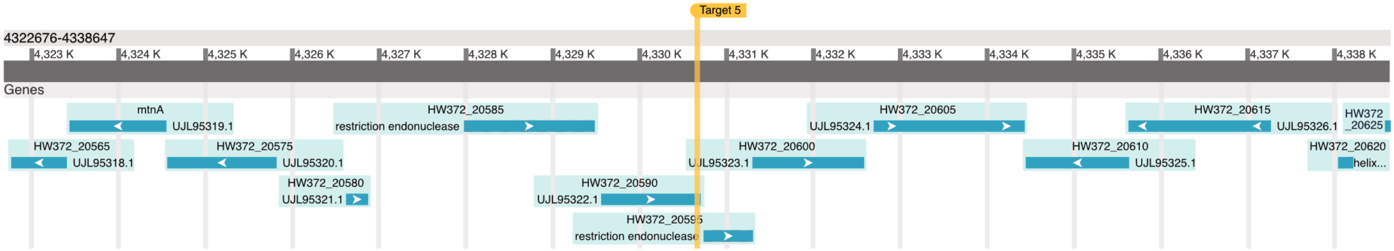

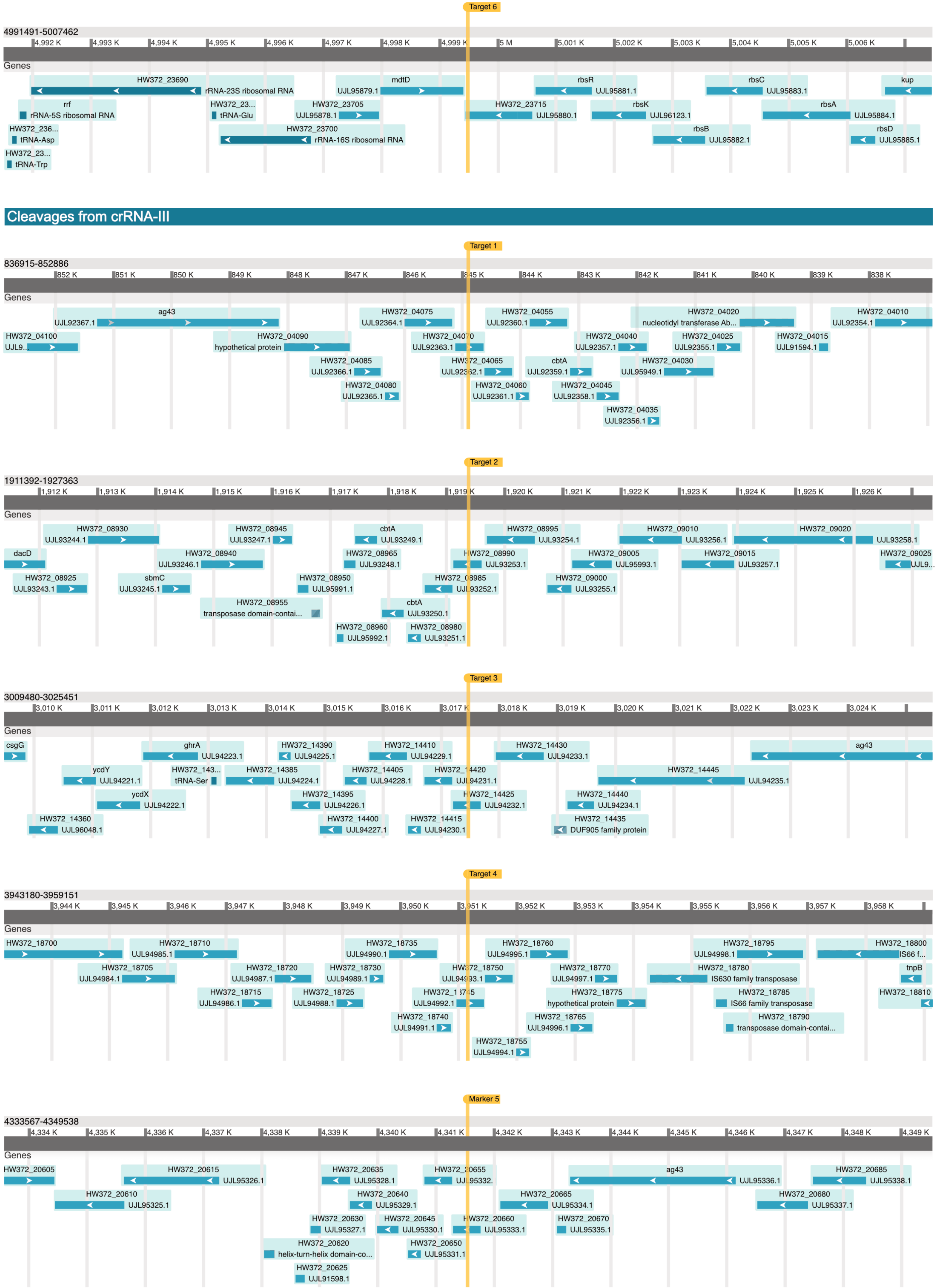

Target 6
4991491-5007462
4,992 K
4,993 K
4,994 K
4,995 K
4,996 K
4,997 K
4,998 K
4,999 K
5 M
5,001 K
5,002 K
5,003 K
5,004 K
5,005 K
5,006 K
Genes
HW372_23690
rRNA-23S ribosomal RNA
mdtD
UJL95879.1
rbsR
UJL95881.1
rbsC
UJL95883.1
kup
rrf
rRNA-5S ribosomal RNA
HW372_23...
tRNA-Glu
HW372_23705
UJL95878.1
HW372_23715
UJL95880.1
rbsK
UJL96123.1
rbsA
UJL95884.1
HW372_236...
tRNA-Asp
HW372_23700
rRNA-16S ribosomal RNA
rbsB
UJL95882.1
rbsD
UJL95885.1
HW372_23...
tRNA-Trp
Cleavages from crRNA-III
Target 1
836915-852886
852 K
851 K
850 K
849 K
848 K
847 K
846 K
845 K
844 K
843 K
842 K
841 K
840 K
839 K
838 K
Genes
ag43
UJL92367.1
HW372_04075
UJL92364.1
HW372_04055
UJL92360.1
HW372_04020
nucleotidyl transferase Ab...
HW372_04010
UJL92354.1
HW372_04100
UJL9...
HW372_04090
hypothetical protein
HW372_04070
UJL92363.1
HW372_04040
UJL92357.1
HW372_04025
UJL92355.1
HW372_04015
UJL91594.1
HW372_04085
UJL92366.1
HW372_04065
UJL92362.1
cbtA
UJL92359.1
HW372_04030
UJL95949.1
HW372_04080
UJL92365.1
HW372_04060
UJL92361.1
HW372_04045
UJL92358.1
HW372_04035
UJL92356.1
Target 2
1911392-1927363
1,912 K
1,913 K
1,914 K
1,915 K
1,916 K
1,917 K
1,918 K
1,919 K
1,920 K
1,921 K
1,922 K
1,923 K
1,924 K
1,925 K
1,926 K
Genes
HW372_08930
UJL93244.1
HW372_08945
UJL93247.1
cbtA
UJL93249.1
HW372_08995
UJL93254.1
HW372_09010
UJL93256.1
HW372_09020
UJL93258.1
dacD
HW372_08940
UJL93246.1
HW372_08965
UJL93248.1
HW372_08990
UJL93253.1
HW372_09005
UJL95993.1
HW372_09015
UJL93257.1
HW372_09025
UJL9...
HW372_08925
UJL93243.1
sbmC
UJL93245.1
HW372_08950
UJL95991.1
HW372_08985
UJL93252.1
HW372_09000
UJL93255.1
HW372_08955
transposase domain-contai...
cbtA
UJL93250.1
HW372_08960
UJL95992.1
HW372_08980
UJL93251.1
Target 3
3009480-3025451
3,010 K
3,011 K
3,012 K
3,013 K
3,014 K
3,015 K
3,016 K
3,017 K
3,018 K
3,019 K
3,020 K
3,021 K
3,022 K
3,023 K
3,024 K
Genes
csgG
ghrA
UJL94223.1
HW372_14390
UJL94225.1
HW372_14410
UJL94229.1
HW372_14430
UJL94233.1
ag43
ycdY
UJL94221.1
HW372_143...
tRNA-Ser
HW372_14385
UJL94224.1
HW372_14405
UJL94228.1
HW372_14420
UJL94231.1
HW372_14445
UJL94235.1
ycdX
UJL94222.1
HW372_14395
UJL94226.1
HW372_14425
UJL94232.1
HW372_14440
UJL94234.1
HW372_14360
UJL96048.1
HW372_14400
UJL94227.1
HW372_14415
UJL94230.1
HW372_14435
DUF905 family protein
Target 4
3943180-3959151
3,944 K
3,945 K
3,946 K
3,947 K
3,948 K
3,949 K
3,950 K
3,951 K
3,952 K
3,953 K
3,954 K
3,955 K
3,956 K
3,957 K
3,958 K
Genes
HW372_18700
HW372_18710
UJL94985.1
HW372_18735
UJL94990.1
HW372_18760
UJL94995.1
HW372_18795
UJL94998.1
HW372_18800
IS66 f...
HW372_18705
UJL94984.1
HW372_18720
UJL94987.1
HW372_18730
UJL94989.1
HW372_18750
UJL94993.1
HW372_18770
UJL94997.1
HW372_18780
IS630 family transposase
tnpB
HW372_18715
UJL94986.1
HW372_18725
UJL94988.1
HW372_18745
UJL94992.1
HW372_18775
hypothetical protein
HW372_18785
IS66 family transposase
HW372_18810
HW372_18740
UJL94991.1
HW372_18765
UJL94996.1
HW372_18790
transposase domain-contai...
HW372_18755
UJL94994.1
Marker 5
4333567-4349538
4,334 K
4,335 K
4,336 K
4,337 K
4,338 K
4,339 K
4,340 K
4,341 K
4,342 K
4,343 K
4,344 K
4,345 K
4,346 K
4,347 K
4,348 K
4,349 K
Genes
HW372_20605
HW372_20615
UJL95326.1
HW372_20635
UJL95328.1
HW372_20655
UJL95332.
ag43
UJL95336.1
HW372_20685
UJL95338.1
HW372_20610
UJL95325.1
HW372_20640
UJL95329.1
HW372_20665
UJL95334.1
HW372_20680
UJL95337.1
HW372_20630
UJL95327.1
HW372_20645
UJL95330.1
HW372_20660
UJL95333.1
HW372_20670
UJL95335.1
HW372_20620
helix-turn-helix domain-co...
HW372_20650
UJL95331.1
HW372_20625
UJL91598.1

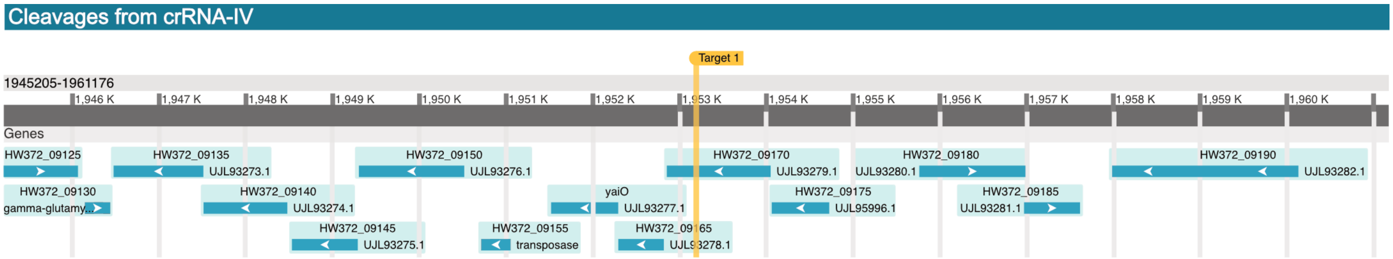
Cleavages from crRNA-IV
Target 1
1945205-1961176
1,946 K
1,947 K
1,948 K
1,949 K
1,950 K
1,951 K
1,952 K
1,953 K
1,954 K
1,955 K
1,956 K
1,957 K
1,958 K
1,959 K
1,960 K
Genes
HW372_09125
HW372_09135
UJL93273.1
HW372_09150
UJL93276.1
HW372_09170
UJL93279.1
HW372_09180
UJL93280.1
HW372_09190
UJL93282.1
HW372_09130
gamma-glutamy...
HW372_09140
UJL93274.1
yaiO
UJL93277.1
HW372_09175
UJL95996.1
HW372_09185
UJL93281.1
HW372_09145
UJL93275.1
HW372_09155
transposase
HW372_09165
UJL93278.1
